# Heparan sulfate regulates amphiregulin signaling towards reparative lung mesenchymal cells during influenza A infection

**DOI:** 10.1101/2024.04.25.591175

**Authors:** Lucas F. Loffredo, Anmol Surpur, Olivia R. Ringham, Fangda Li, Kenia de los Santos-Alexis, Nicholas Arpaia

## Abstract

Amphiregulin (Areg), a growth factor produced by regulatory T (Treg) cells to facilitate tissue repair/regeneration, contains a heparan sulfate (HS) binding domain. How HS, a highly sulfated glycan subtype that alters growth factor signaling, influences Areg repair/regeneration functions is unclear. Here we report that inhibition of HS in various cell lines and primary lung mesenchymal cells (LMC) qualitatively alters downstream signaling and highlights the existence of HS-dependent vs. -independent Areg transcriptional signatures. Utilizing a panel of cell lines with targeted deletions in HS synthesis–related genes, we found that the presence of the glypican family of heparan sulfate proteoglycans is critical for Areg signaling and confirmed this dependency in primary LMC by siRNA-mediated knockdown. Furthermore, in the context of influenza A (IAV) infection *in vivo*, we found that an Areg-responsive subset of reparative LMC upregulate glypican-4 and HS. Conditional deletion of HS primarily within this LMC subset resulted in reduced blood oxygen saturation following infection with IAV, with no changes in viral load. Finally, we found that co-culture of HS-knockout LMC with IAV-induced Treg cells results in reduced LMC responses. Collectively, this study reveals the essentiality of HS on a specific lung mesenchymal population as a mediator of Treg cell–derived Areg reparative signaling during IAV infection.

## Introduction

In order to engage in their myriad functions, cells of the immune system must migrate through, work within, and communicate with dense tissue networks of non-immune cells. While the implications of these interactions are well-studied in the context of anti-pathogen responses, our knowledge of the role these interactions play in regulating tissue homeostasis and repair remains far more limited.

One mechanism by which immune cells mediate tissue regulatory changes is via production of amphiregulin (Areg). Areg is a growth factor of the epidermal growth factor receptor (EGFR) ligand family that has been identified as an immune-derived mediator of repair in multiple tissue damage and disease contexts [1]. Previous reports have demonstrated that lung regulatory T (Treg) cells produce Areg in the context of influenza A virus (IAV) infection in mice, and that genetic deletion of *Areg* specifically in Treg cells impairs proper recovery of blood oxygen saturation (SpO_2_), a feature found to be independent of the canonical role Treg cells play in immunosuppression [2]. While these tissue effects were initially thought to be the result of Areg signaling directly to epithelial cells, further work from our group demonstrated that a distinct subset of lung mesenchymal cells (LMC) are in fact the critical Areg-sensing intermediate in this context [3]. This LMC population showed high EGFR expression, unique responsiveness to Areg, and spatial localization near areas of alveolar damage, characteristics that endow this subset with the ability to send reparative signals to epithelial cells in response to Treg cell–derived Areg. We and others have referred to this cell type as “*Col14a1*^+^ lung mesenchymal cells” (abbreviated here as “Col14-LMC”) [4], based on high expression of *Col14a1* (encoding Type XIV collagen); transcriptionally similar LMC populations have been referred to in other publications as mesenchymal alveolar niche cells (MANC) or adventitial fibroblasts [5-8]. Our previous work also included comparisons to the other two most sizable subsets of mouse LMC, marked by expression of *Col13a1* (“Col13-LMC”) or *Hhip* (“Hhip-LMC”), alternatively referred to in the literature as alveolar fibroblasts and peribronchial fibroblasts, respectively [7]; these LMC populations expressed lower amounts of EGFR and were less responsive to Areg stimulation.

Heparan sulfate (HS) is a type of glycosaminoglycan (GAG) that is produced and presented by tissue cells (see **Fig. 1A** for visual summary of HS biology to accompany this description). Among the diverse types of GAGs, HS has received increased attention due to its ability to harbor the greatest amount of sulfate groups, giving it a highly negative charge [9] and allowing it to affect the signaling of a diverse array of positively charged HS binding proteins (HSBPs) [10]. HS is produced and presented in a protein-bound form, either on cell surface proteins (syndecans, which have a transmembrane domain, or glypicans, which are glycosylphosphatidylinositol [GPI]-anchored), or on secreted extracellular matrix (ECM) proteins [9]. HS is initially synthesized in the Golgi apparatus as an unsulfated polysaccharide chain consisting largely of repeating glucuronic acid and N-acetylglucosamine disaccharide subunits; this chain is then altered by sulfate group addition during Golgi apparatus trafficking by various sulfotransferase enzymes, which impart its negative charge [9]. Significant heterogeneity exists along each individual HS chain, with different areas containing low or high levels of sulfation [9]. From past work on HS, the paradigm has emerged that HS is ubiquitously expressed on all tissue cells [11]; however, most reports studying HS in tissue environments do not explicitly evaluate levels of HS on various tissue cell types.

**Figure 1.**
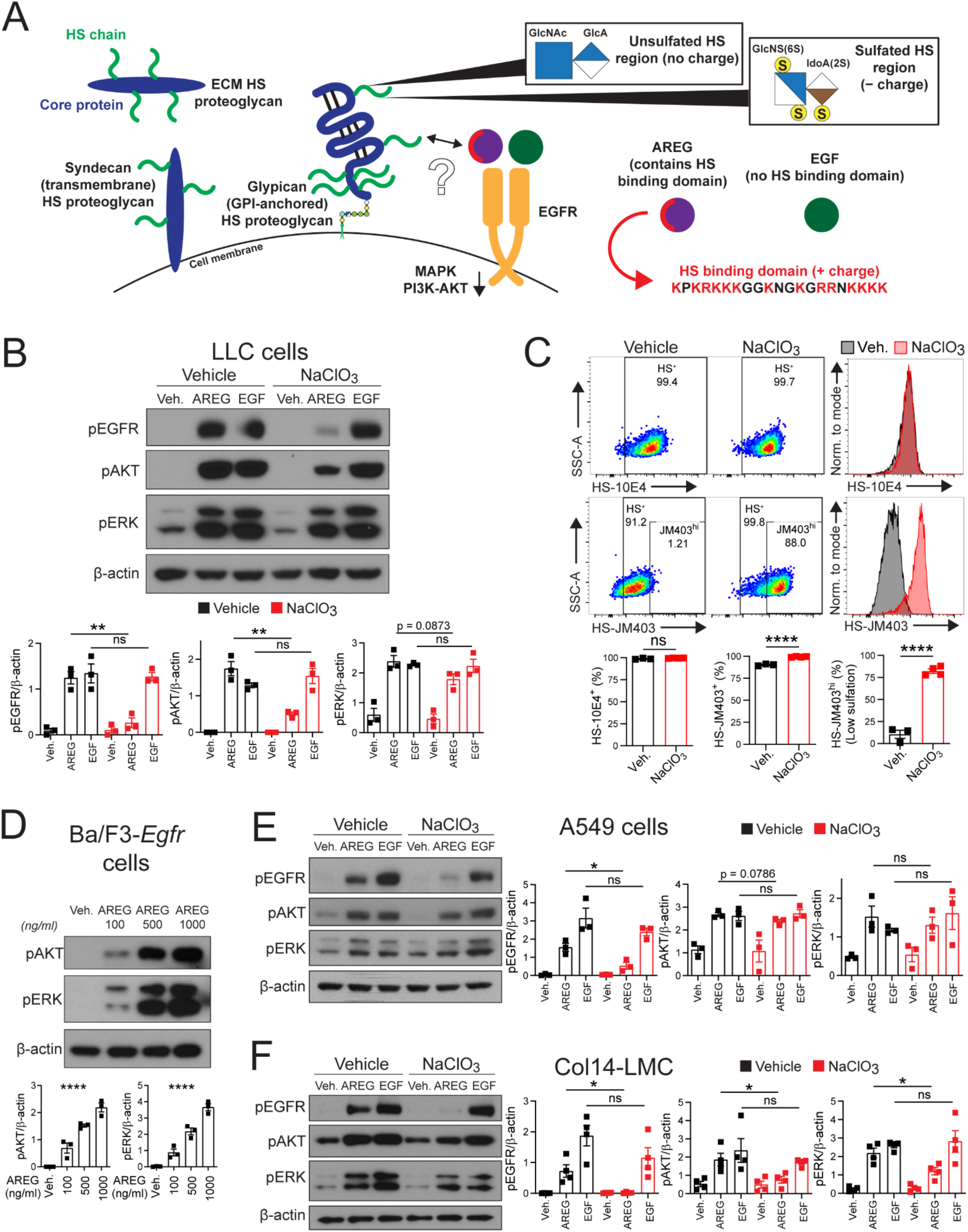
Areg signaling is altered but not abrogated in the context of HS inhibition. **(A)** Diagrammatic summary of different types of HS-presenting proteoglycans, heterogeneity in sulfation along a single HS chain, and EGFR ligand signaling interactions with HS. HS: heparan sulfate; ECM: extracellular matrix; GlcNAc: N-acetylglucosamine; GlcA: glucuronic acid; GlcNS(6S): N-sulfoglucosamine (with 6-O-sulfation); IdoA(2S): iduronic acid (with 2-O-sulfation); S in yellow circle: sulfate group; EGFR: epidermal growth factor receptor; AREG: amphiregulin; EGF: epidermal growth factor. **(B)** Western blotting for phospho-EGFR (Y1068), phospho-AKT (S473), phospho-ERK (T202/Y204), and β-actin of vehicle or sodium chlorate (NaClO_3_)-treated (16-18h) LLC cells, stimulated for 15 min. with vehicle, murine AREG (500 ng/ml), or murine EGF (100 ng/ml). Representative western blots shown. n=3 per condition, graphs contain all values from 3 separate experiments. **(C)** Flow cytometry using HS-directed antibodies 10E4 or JM403 on LLC cells treated with vehicle or NaClO_3_ (16-18h). Representative flow cytometry plots shown. Gating based on fluorescence-minus-one (FMO) controls. Percent staining positive displayed in plots. n=3-4 per condition, graphs contain all values from 2 separate experiments. **(D)** Western blotting for phospho-AKT (S473), phospho-ERK (T202/Y204), and β-actin of vehicle or murine AREG-stimulated (at various concentrations) (15 min.) Ba/F3-*Egfr* cells. Representative western blots shown. n=3 per condition, graphs contain all values from 3 separate experiments. **(E)** Western blotting for phospho-EGFR (Y1068), phospho-AKT (S473), phospho-ERK (T202/Y204), and β-actin of vehicle or NaClO_3_-treated (16-18h) A549 cells, stimulated for 15 min. with vehicle, human AREG (500 ng/ml) or EGF (100 ng/ml). Representative western blots shown. n=3 per condition, graphs contain all values from 3 separate experiments. **(F)** Western blotting for phospho-EGFR (Y1068), phospho-AKT (S473) phospho-ERK (T202/Y204), and β-actin of vehicle or NaClO_3_-treated (16-18h) Col14-LMC, stimulated for 15 min. with vehicle, murine AREG (500 ng/ml), or EGF (100 ng/ml). Col14-LMC were gated/sorted as shown in Figure S2 (negative bead enriched), then at 24h post-plating non-adherent cells were washed away/media changed, with subsequent vehicle or NaClO_3_ treatment. Representative western blots shown. n=4 per condition, graphs contain all values from 4 separate experiments. Standard error displayed on graphs; n.s: not significant, *: 0.01<p<0.05, **: 0.001<p<0.01, ***: 0.0001<p<0.001, ****: p<0.0001.

Unlike the canonical EGFR ligand family member epidermal growth factor (EGF), Areg contains a HS binding domain. While early reports found that Areg is dependent on HS for signaling *in vitro* [12, 13], few studies have investigated interactions between Areg and HS since this initial work. Interestingly, while there are numerous cellular sources for Areg (e.g., epithelial cells), only Areg from certain immune cells in specific disease contexts effectively supports tissue repair, implying a highly spatially localized component for its tissue regulatory roles. We reasoned that due to this apparent spatial specificity of action for Areg, and since HS expression varies greatly among tissue cell types in settings of inflammation, there may be unappreciated aspects of Areg–HS interactions in these processes that merit further investigation.

In this report, we sought to broaden the investigation of Areg–HS interactions by using chemical and genetic inhibition of HS *in vitro*, in order to analyze downstream signaling, transcriptional effects, and molecular components of this relationship. We furthered these inquiries to an *in vivo* tissue context using models of lung damage and novel conditional HS deletion mice targeting a specific subpopulation of LMC. Furthermore, we found that both a specific core protein for HS presentation (glypican-4) and HS itself are uniquely upregulated on Areg target cells in the lung (Col14-LMC) during IAV infection, endowing them with heightened ability to respond to reparative Treg cells. Our findings demonstrate a critical role for HS on a distinct subset of LMC, which is upregulated and utilized in order to respond to immune cell–derived Areg and enact proper tissue repair modalities following IAV-induced damage.

## Results

### Areg signaling is altered but not abrogated in the context of HS inhibition

To confirm and extend earlier observations regarding dependency of the Areg HS binding domain for signaling **(Fig. 1A)** [12, 13], we performed experiments using the Lewis lung carcinoma (LLC) mouse epitheloid cancer cell line, which has high expression of HS and EGFR [14, 15], using known HS signaling inhibitors. Notably, while Areg signaling dependency on HS was previously only shown using an indirect readout of DNA synthesis, we chose to test this molecularly by assessing phosphorylation of EGFR via western blotting in response to ligand stimulation. EGF, which lacks an HS binding domain, was used as a control. The dependency on HS for Areg-, but not EGF-signaling, was shown using three HS-altering treatment methods: *(1)* sodium chlorate (NaClO_3_), which prevents formation of sulfate donors in the cytoplasm, thus reducing sulfation in displayed HS [12] **(Fig. 1B)**, *(2)* heparinase I/III, which enzymatically cleaves HS from the surface of cells [12] **(Fig. S1A)**, and *(3)* surfen, a positively charged small molecule antagonist of HS which functions by binding up negatively charged sulfated regions of HS to make them refractory to HSBP engagement [16] **(Fig. S1B)**.

To assess the effects of these HS-altering treatments on LLC cell surface-bound HS, we used two established antibodies for HS assessment: 10E4, which targets highly sulfated regions on HS [17], and JM403, which targets regions of low sulfation on HS [18]. As visualized by 10E4 staining, NaClO_3_ treatment did not prevent the production and presentation of HS by LLC cells; in fact, enough sulfated HS regions are maintained on cells in this context to prevent any reduction in 10E4 staining **(Fig. 1C)**. However, staining with JM403 shows a full shift in the profile of LLC cells from a JM403^lo^ to JM403^hi^ phenotype upon NaClO_3_ treatment **(Fig. 1C)**. This is an indication that low-sulfation regions of HS predominate upon treatment with NaClO_3_. Heparinase I/III treatment of LLC cells resulted in reduced staining for 10E4 and JM403 **(Fig. S1C)**. Surfen treatment did not alter staining for 10E4 or JM403 on LLC cells **(Fig. S1D)**.

Downstream signaling via EGFR occurs through the MAPK and the AKT-mTOR pathways [19]. We evaluated signaling via these pathways from Areg and EGF treatment with or without HS sulfation inhibition (NaClO_3_ treatment) using western blotting, expecting this to mirror the full inhibition seen for EGFR phosphorylation. Instead, we found that in the context of HS sulfation inhibition (NaClO_3_) in Areg-stimulated cells, ERK phosphorylation (MAPK pathway) is maintained at similar levels, while AKT phosphorylation is significantly reduced but still present, compared to vehicle controls **(Fig. 1B)**. Prior research investigating Areg signaling dependence on HS has implied that without HS, Areg is fully deficient in signaling; our findings run counter to this, indicating that there are HS-dependent and HS-independent modalities of Areg signaling.

To eliminate the possibility that residual MAPK and AKT signaling in the setting of NaClO_3_ treatment is due to impartial HS inhibition, we tested the ability of Areg to signal in Ba/F3 cells, a murine pro-B cell line reported to completely lack HS [20] – which we confirmed using HS-targeting antibodies **(Fig. S1E)**. This cell line also lacks expression of EGFR [21]; thus, we transiently transfected a construct encoding mouse *Egfr* into these cells, and confirmed its signaling capability via treatment with EGF **(Fig. S1F)**. Upon treatment with Areg, we saw a dose-dependent response in downstream ERK and AKT phosphorylation in these cells **(Fig. 1D)** (EGFR phosphorylation was not able to be assessed in this cell type). The dose dependency of this response indicates that Areg is binding and inducing *bona fide* activation of its receptor, in a context completely devoid of HS.

An additional method we utilized to test the effects of HS on Areg signaling was to use a “pre-binding” strategy. Recombinant Areg was first incubated with free HS to block available HS binding sites prior to applying this mixture to LLC cells, so that any residual signaling toward EGFR must occur independently of this binding domain. Using this strategy, we found that EGFR phosphorylation and AKT phosphorylation were significantly reduced, but ERK phosphorylation remained fully intact **(Fig. S1G)**. These results further confirmed that the HS binding domain alters Areg signaling, primarily affecting the AKT-mTOR pathway rather than the MAPK pathway.

To test that this is also relevant for human Areg, we performed select experiments on A549 cells, a human adenocarcinoma alveolar epithelial cell line with ubiquitous expression of HS and high expression of EGFR [22, 23]. NaClO_3_ treatment largely recapitulates our previous results seen in LLC cells upon Areg and EGF stimulation – EGFR phosphorylation is significantly reduced, AKT phosphorylation shows a trend towards reduction but is still present, and ERK phosphorylation maintains full signaling potential **(Fig. 1E)**. Similarly, NaClO_3_ treatment of A549 cells promotes a wholesale shift of JM403 staining from a low- to high-staining profile **(Fig. S1H)**. Thus, the HS-dependent vs. -independent signaling modalities of Areg are also present in the human context.

Lastly, we investigated whether a similar signaling pattern is present in primary cells relevant for Areg signaling; we chose to use Col14-LMC for this cell type, given their EGFR^hi^ phenotype and reparative interaction with Treg cell–derived Areg described in our previous publication (Kaiser 2023). In our prior study, we demonstrated the ability to isolate and culture the major subsets of lung mesenchymal cells described above (Col14-LMC, Col13-LMC, and Hhip-LMC), via flow cytometric sorting from enriched lung mesenchyme **(Fig. S2)** [3]. Upon sorting and culturing of Col14-LMC, we found that stimulation with Areg in the presence of NaClO_3_ treatment resulted in significantly decreased EGFR and AKT phosphorylation, similar to that seen in LLC cells **(Fig. 1F)**. For MAPK pathway signaling, a significant decrease in ERK phosphorylation was observed for Areg stimulation in the presence of NaClO_3_; however, an increase in ERK phosphorylation was apparent when compared to NaClO_3_ treatment alone **(Fig. 1F)**, confirming that Col14-LMC are able to undergo HS-independent signaling as described above for LLC. We also observed a similar surface HS pattern as was observed in LLC when Col14-LMC were stained with HS-targeted antibody JM403 in the presence of NaClO_3_ treatment (shift from JM403^lo^ to JM403^hi^) **(Fig. S1I)**. Based on these observations, we concluded that the HS-dependent and -independent signaling modalities are preserved in this Areg signaling–relevant primary cell type.

### Transcriptional profile of HS-dependent vs. -independent Areg signaling in Col14-LMC

We reasoned that HS-dependent vs. -independent signaling may impact the transcriptional signature of target cells. To this end, we performed bulk RNA-seq on vehicle-treated/Areg-stimulated vs. NaClO_3_-treated/Areg-stimulated Col14-LMC. Areg treatment was administered at three different concentrations; this was done to allow for comparison between similarly sized gene signatures for HS-unaltered vs. HS sulfation-inhibited conditions, in the case that the latter resulted in overall lessened gene expression changes in response to Areg stimulation. Observing total numbers of up/downregulated differentially expressed genes (DEGs) in these conditions **(Fig. 2A)**, this was indeed the case, with NaClO_3_ Col14-LMC showing lower DEGs compared to vehicle treated at each concentration of Areg tested. However, HS sulfation-inhibited Col14-LMC still exhibit a dose-responsive increase in DEGs in response to increasing Areg concentrations, with an appreciable gene signature of 2504 total DEGs at the highest concentration, confirming our previous observation that Areg is able to signal in the absence of proper HS engagement. Principle component analysis of full gene expression profiles identified Areg responsiveness as the primary driver of separation between samples, with little separation seen on the secondary axis between vehicle and NaClO_3_-exposed samples, consistent with an evident but smaller (in comparison to Areg treatment) effect of NaClO_3_ exposure at baseline (1059 DEGs) **(Fig. 2B)**.

**Figure 2.**
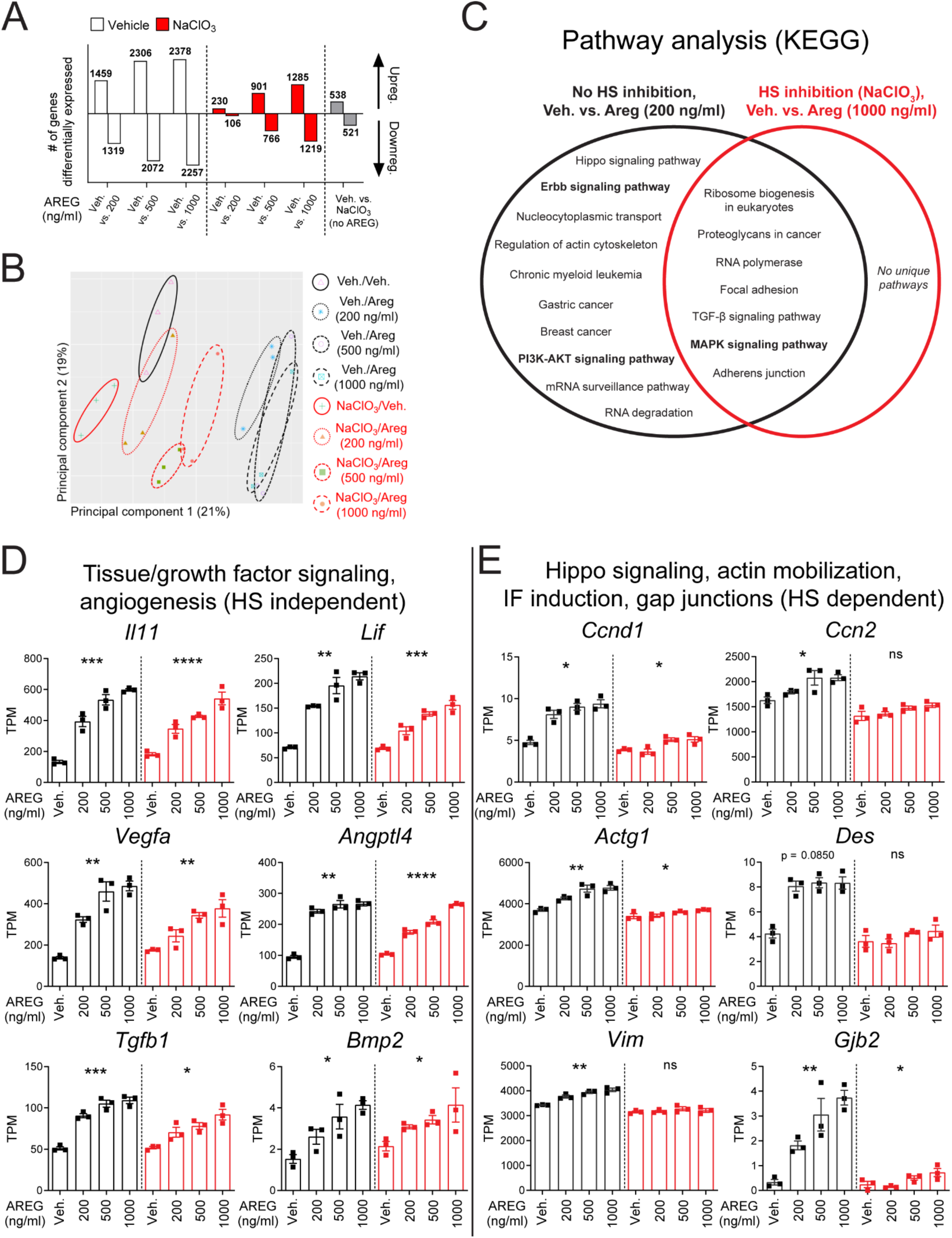
Transcriptional profile of HS-dependent vs. -independent Areg signaling in Col14-LMC. Bulk RNA-seq of Col14-LMC treated with vehicle or NaClO_3_ (16-18h), then stimulated with varying concentrations of murine AREG for 4h. Col14-LMC were gated/sorted as shown in Figure S2 (negative bead enriched), then at 24h post-plating non-adherent cells were washed away/media changed, with subsequent vehicle or NaClO_3_ treatment (16-18h) prior to AREG stimulation. **(A)** Significant differentially expressed genes determined by DESeq2 (FDR-adjusted P-value < 0.05, no FC cutoff), for comparisons indicated on x-axis. **(B)** Principal components analysis of all samples from bulk RNA-seq of Col14-LMCs treated with vehicle or NaClO_3_ (16-18h) and stimulated with varying concentrations of murine AREG. **(C)** Pathway analysis (gProfiler, KEGG pathways) on upregulated genes from RNA-seq, from the “No HS inhibition, vehicle vs. AREG (200 ng/ml)” comparison (1459 DEGs) (black circle), and the “HS inhibition, vehicle vs. AREG (1000 ng/ml)” comparison (1285 DEGs) (red circle). Certain pathways highlighted in Results section boldened. **(D–E)** Transcript per million (TPM) values from RNA-seq for representative genes from select functional pathways; significance across samples in each group (n=3) calculated by Kruskal-Wallis test. IF: intermediate filament. Standard error displayed on graphs; n.s: not significant, *: 0.01<p<0.05, **: 0.001<p<0.01, ***: 0.0001<p<0.001, ****: p<0.0001.

In order to compare HS-intact and HS sulfation-inhibited cells at similar overall numbers of identified DEGs, we performed pathway analysis (KEGG pathways) on upregulated genes from the lowest Areg concentration for vehicle-treated Col14-LMC (200 ng/ml; 1459 upreg. DEGs) and the highest Areg concentration for NaClO_3_-treated Col14-LMC (1000 ng/ml; 1285 upreg. DEGs). Significantly upregulated pathways for each are summarized in **Fig. 2C**; pathways upregulated in both contexts (middle of Venn diagram) are HS-independent (i.e., they are upregulated regardless of HS sulfation inhibition), while pathways upregulated only in the HS-intact context (left of Venn diagram) are HS-dependent. There were no significantly upregulated pathways represented in the HS sulfation inhibition context that were not represented in the HS-intact context. Strikingly, mirroring our previous results at the signaling level, the MAPK signaling pathway is significantly upregulated in both contexts (HS-independent), while the ERBB signaling pathway and the PI3K-AKT signaling pathway are only upregulated in the HS-intact context (HS-dependent).

Genes involved in tissue/growth factor signaling and angiogenesis (e.g., *Il11*, *Lif*, *Vegfa*, *Angtl4*, *Tgfb1*, *Bmp2*) show significant upregulation across groups in both HS-intact and sulfation-inhibited conditions, with similar or only slightly reduced levels of induction in the latter **(Fig. 2D)**, demonstrating that these tissue modalities appear to be largely independent of HS involvement. Other pathways were only substantially upregulated across groups in the HS-intact scenario (HS-dependent), including genes related to the Hippo pathway, actin mobilization, and intermediate filament induction (e.g., *Ccnd1*, *Ccn2*, *Actg1*, *Des*, *Vim*) (**Fig. 2E**). These modalities have the potential to regulate cell fate, cell polarity, and movement in response to an activating stimulus [24]. Interestingly, *Gjb2*, the gene encoding connexin-26, shows this pattern as well; this gene is critical for proper formation of gap junctions [25], which may point to a role in fostering intercellular communication as an additional HS-dependent function.

### HS-related gene knockout cell lines identify glypican as a critical HS core protein for Areg signaling

We next sought to understand the specific characteristics of HS that confer its ability to interact with Areg. The genes encoding mediators of HS synthesis and modification are summarized in **Figure S3** [9]. To ascertain which of these HS-related genes are critical for Areg signaling, we created a panel of CRISPR-Cas9-mediated knockout (KO) LLC cell lines. We began by querying a publicly available RNA-seq dataset of WT LLC cells to get a baseline layout of HS-related gene expression in this cell line **(Fig. S4A)** [26]. We chose to include enzymes involved in HS extension and sulfation, as well as certain core proteins that harbor HS **(Fig. 3A)**, while avoiding genes involved in HS chain initiation due to their myriad roles in forming other types of glycans and additional cellular processes [9]. Two separate single cell–derived clonal lines were generated for each targeted gene, with the exception of *Sdc1*, where we were only able to generate one subline. For all KO sublines, we confirmed a reduction in expression at the mRNA level **(Fig. S4B)**, and modification of HS (for enzymes) or reduction in core protein levels (as discussed below).

**Figure 3.**
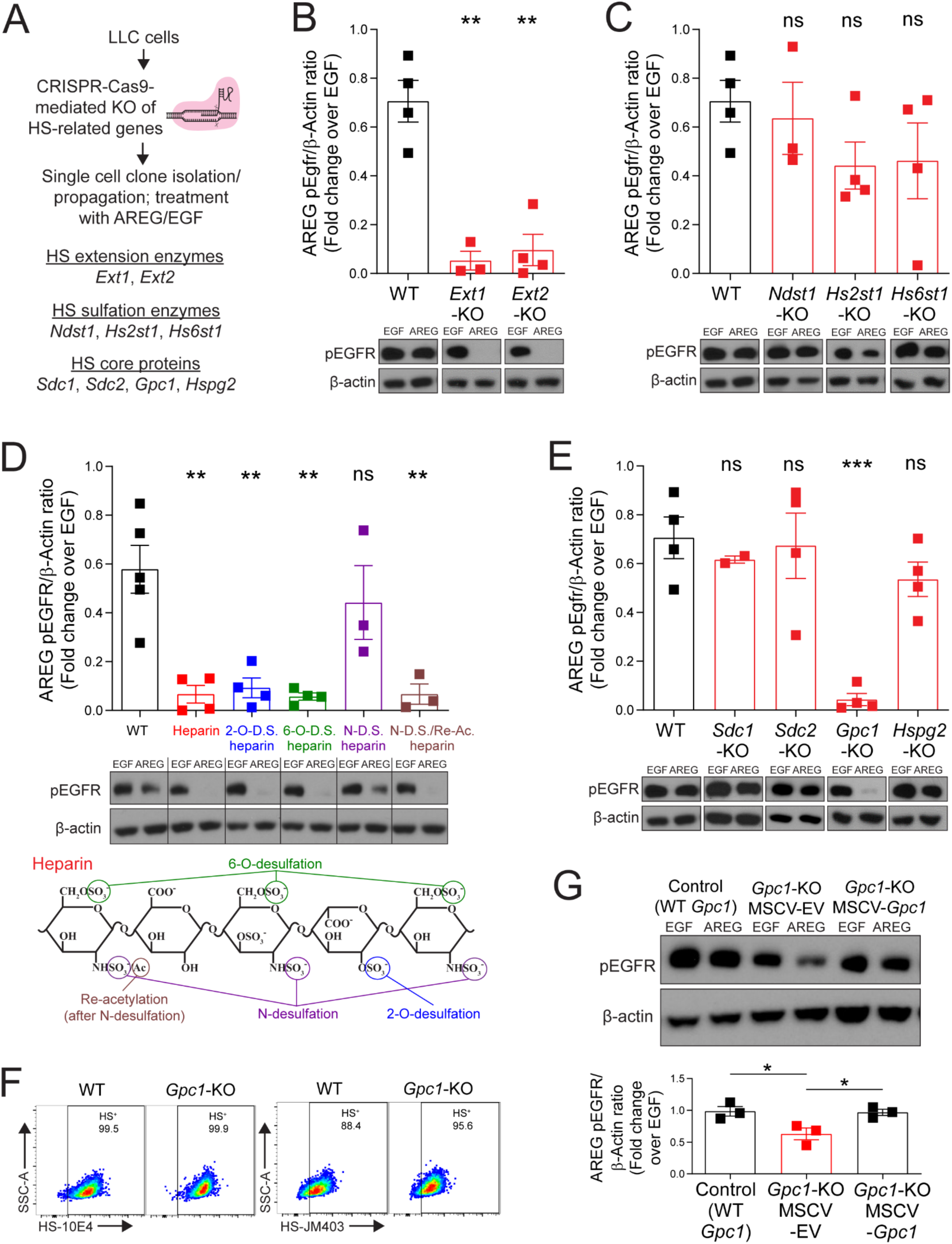
HS-related gene knockout cell lines identify glypican as a critical HS core protein for Areg signaling. **(A)** Experimental schematic for generation of knockout (KO) panel of HS-related genes by CRISPR-Cas9 in LLC cells. **(B)** Western blotting for phospho-EGFR (Y1068) and β-actin of WT and *Ext1*- or *Ext2*-KO LLC cell sublines, stimulated for 15 min. with murine AREG (500 ng/ml) or EGF (100 ng/ml). AREG phosphorylation level quantification was done by adjusting to EGF controls. Representative western blots shown. n=3-4 per gene, graph contains all values from 3-4 separate experiments (across 2 clonal sublines per gene). **(C)** Western blotting for phospho-EGFR (Y1068) and β-actin of WT and *Ndst1*-, *Hs2st1*-, or *Hs6st1*-KO LLC cell sublines, stimulated for 15 min. with murine AREG (500 ng/ml) or EGF (100 ng/ml). AREG phosphorylation level quantification was done by adjusting to EGF controls. WT control blots are the same as those used in B, as these experiments were ran and analyzed on the same blots. Representative western blots shown. n=3-4 per gene, graph contains all values from 3-4 separate experiments (across 2 clonal sublines per gene). **(D)** Western blotting for phospho-EGFR (Y1068) and β-actin of WT LLC stimulated for 15 min. with murine AREG (500 ng/ml) or EGF (100 ng/ml) that had been pretreated for 15 min. with vehicle or heparin (WT or chemically desulfated at specific sites). AREG phosphorylation level quantification was done by adjusting to EGF controls. Representative western blots shown. n=3-4 per heparin variant, graph contains all values from 3-4 separate experiments. Schematic included of targeted desulfation/reacetylation sites in differentially desulfated heparin variants. **(E)** Western blotting for phospho-EGFR (Y1068) and β-actin of WT and *Sdc1*-, *Sdc2*-, *Gpc1*-, or *Hspg2*-KO LLC cell sublines, stimulated for 15 min. with murine AREG (500 ng/ml) or EGF (100 ng/ml). AREG phosphorylation level quantification was done by adjusting to EGF controls. WT control data are the same as those used in B, as these experiments were ran and analyzed on the same blots. Representative western blots shown. n=2-4 per gene, graph contains all values from 2-4 separate experiments (across 2 clonal sublines per gene, for all except *Sdc1* [only 1 clone]). **(F)** Flow cytometry using HS-directed antibodies 10E4 and JM403 on WT or *Gpc1*-KO LLC. Representative plots shown from 3 separate experiments. **(G)** Western blotting for phospho-EGFR (Y1068) and β-actin of CRISPR-Cas9 control (WT *Gpc1*), *Gpc1*-KO LLC transduced with empty vector (EV) MSCV retrovirus (MSCV-EV), or *Gpc1*-KO LLC transduced with MSCV retrovirus with *Gpc1* mRNA (MSCV-*Gpc1*), stimulated for 15 min. with murine AREG (500 ng/ml) or EGF (100 ng/ml). AREG phosphorylation level quantification was done by adjusting to EGF controls. Representative western blots shown. n=3 per condition, graph contains all values from 3 separate experiments. Standard error displayed on graphs; n.s: not significant, *: 0.01<p<0.05, **: 0.001<p<0.01, ***: 0.0001<p<0.001, ****: p<0.0001.

*Ext1* and *Ext2* are non-redundantly responsible for HS chain extension (and are not involved in formation of other glycans); thus, we anticipated that KO of these genes would result in full HS ablation. Accordingly, upon knockout of *Ext1* or *Ext2* in LLC, we saw an elimination of staining for HS-directed antibodies 10E4 and JM403 **(Fig. S4C)**. As expected, *Ext1*- and *Ext2*-KO sublines show a profound reduction in Areg signaling induction, as quantified by EGFR phosphorylation, compared to WT controls **(Fig. 3B)**.

To alter sulfation of HS, we chose to delete *Ndst1*, *Hs2st1*, and *Hs6st1* – highly expressed representative genes of the N-sulfotransferase, 2-O-sulfotransferase, and 6-O-sulfotransferase families responsible for sulfation of different sites on HS. While western blots attempting to demonstrate protein-level depletion in sulfation enzyme KO sublines were unclear, significant alterations were viewed in the HS profile of these sublines using 10E4 and JM403 antibodies: *Ndst1*-KO sublines show a reduction in 10E4 staining and an increase in JM403 staining; *Hs2st1*-KO sublines show a reduction in JM403 staining; and *Hs6st1*-KO sublines show an increase in 10E4 and JM403 staining **(Fig. S4D)**. Based on prior paradigms in the field indicating that sulfation state of HS is the determinative factor in its ability to bind HS-binding proteins [10], we expected that KO of the sulfotransferase enzymes would be effective in altering Areg signaling; however, we found that none of the sulfotransferase enzyme KO sublines exhibited reduced Areg signaling compared to WT controls **(Fig. 3C)**.

To determine why deletions in sulfotransferase genes did not affect Areg signaling, we engaged in another line of experimentation to test the ability of Areg to bind to HS with altered sulfation patterns. Heparin is a highly sulfated subtype of HS that is widely used as a clinical anticoagulant. Using a similar Areg pre-binding strategy as in **Figure S1H**, we tested the ability of different heparin variants, chemically modified to lack sulfation at certain sites, to bind Areg and inhibit EGFR signaling. As expected, we found that Areg pre-bound to heparin was unable to induce signaling **(Fig. 3D)**. Of the differentially desulfated heparin variants, only N-desulfated heparin showed reduced binding to Areg, as demonstrated by similar ability of Areg to induce signaling when pre-incubated with this variant compared to vehicle-exposed Areg **(Fig. 3D)**. This finding could be taken as evidence that sulfation at the N-site is important for Areg binding to heparin and/or HS; however, in light of the fact that Areg signaling is inhibited by pre-incubation with a reacetylated form of N-desulfated heparin, we believe it more likely that a residual positive charge on N-desulfated heparin (prior to re-acetylation) is leading to repulsive ionic forces that are weakening Areg binding to this variant. Thus, when taken together with the results from our panel of sulfotransferase KO clones **(Fig. 3C),** these data indicate that Areg does not show a strong preference for specific sulfation modifications in its binding to HS, rather the net negative charge of heparan sulfate mediates its interaction with Areg.

With regards to cell surface core proteins, while analysis of publicly available RNA-seq datasets indicated expression of *Sdc1*, *Sdc2*, *Sdc4*, *Gpc1*, and *Gpc6* by LLC, staining with antibodies directed towards these proteins showed only appreciable expression of Sdc1, Sdc2, and Gpc1 **(Fig. S4E)**; thus, we chose to explore the contribution of each of these core proteins using our CRISPR-mediated knockout strategy. Knockouts of secreted ECM (*Hspg2*) and cell surface–localized (*Sdc1*, *Sdc2*, *Gpc1*) HS core proteins were confirmed at the protein level via flow cytometry **(Fig. S4F)**. Surprisingly, despite the concept in the HS field that core proteins are generally not a determinative factor in HSBP signaling, we found that in *Gpc1*-KO sublines (but not *Sdc1*-KO, *Sdc2*-KO, or *Hspg2*-KO), Areg signaling was significantly reduced compared to EGF controls **(Fig. 3E)**. Strikingly, despite the lack of Gpc1 protein in KO cell lines, these cells showed fully intact presentation of HS on their surface using 10E4 and JM403 antibodies, likely presented by other cell surface–localized HS core proteins **(Fig. 3F)**; thus, this reduction in Areg signaling in *Gpc1*-KO sublines is not the result of global defects in HS presentation. To confirm the specificity of this signaling reduction, we performed rescue experiments using *Gpc1*-KO cells, reinstating expression via transduction with a retroviral vector encoding *Gpc1* **(Fig. S4G)**. Upon rescue of *Gpc1* expression, Areg signaling was fully restored **(Fig. 3G)**. From these experiments, we conclude that glypicans (i.e., GPI-anchored HS-presenting core proteins) are critical for proper Areg interaction with HS in a signaling context.

### Glypicans are critical for proper Areg signaling in primary Areg-responsive cells and are upregulated specifically on Col14-LMC in the context of viral lung infection

We next sought to determine if specific HS-presenting core proteins (glypicans) also serve as critical determinants of Areg signaling in LMC. We first queried our previously published bulk RNA-seq dataset and a publicly available scRNA-seq dataset to inform which cell surface core proteins are present in murine LMC [3, 5] **(Fig. S5A, S5B)**. Across bulk RNA-seq and scRNA-seq datasets, various syndecans (*Sdc1*, *Sdc2*, *Sdc3*, *Sdc4*) and glypicans (*Gpc1*, *Gpc3*, *Gpc4*, *Gpc6*) show appreciable transcriptional expression; however, at the protein level, Sdc2 was the most highly expressed syndecan and Gpc4 was the only glypican for which we could detect appreciable expression levels **(Fig. 4A)**.

**Figure 4.**
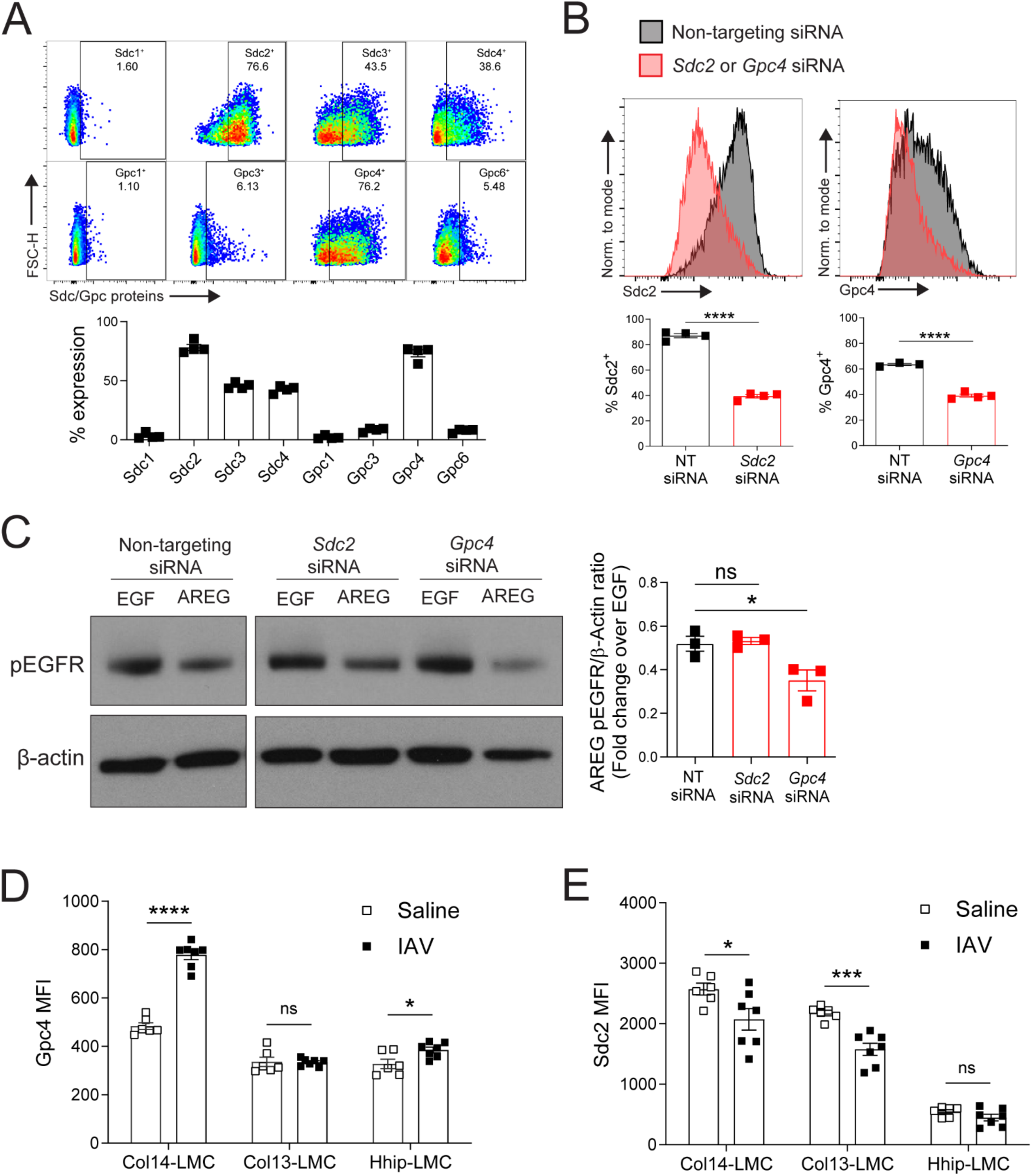
Glypicans are critical for proper Areg signaling in primary Areg-responsive cells and are upregulated specifically on Col14-LMC in the context of viral lung infection. **(A)** Flow cytometry using antibodies targeting Sdc1, Sdc3, Sdc4, Gpc1, Gpc3, or Gpc6 on cultured bulk LMC (negative bead enrichment for CD45^-^CD31^-^Epcam^-^ cells, plated without sorting [see Figure S2]; non-adherent cells washed away/media changed at 16-18h, then cultured for 48h). Gated on CD45^-^ CD31^-^Epcam^-^Pdgfra^+^ cells post-culture. Representative flow cytometry plots shown. Gating based on FMO controls. Percent staining positive displayed in plots. n=4 per target, graph contains all values from 2 separate experiments. **(B)** Flow cytometry using antibodies targeting Sdc2 or Gpc4 on cultured bulk LMC, treated with non-targeting (NT) siRNA or siRNA targeting either *Sdc2* or *Gpc4* (negative bead enrichment as in A; non-adherent cells washed away/media changed after 16-18h, then treated with 25-50 nM siRNA for 48h). Gated on CD45^-^CD31^-^Epcam^-^Pdgfra^+^ cells post-culture. Representative flow cytometry plots shown. n=3-4 per target, graphs contain all values from 2-3 separate experiments. **(C)** Western blotting for phospho-EGFR (Y1068) and β-actin of bulk LMC treated with NT siRNA, *Sdc2* siRNA, or *Gpc4* siRNA as in B (50 nM, 48h), then stimulated for 15 min. with murine AREG (500 ng/ml) or EGF (100 ng/ml). AREG phosphorylation level quantification was done by adjusting to EGF phosphorylation controls. Representative western blots shown. n=3 per condition, graph contains all values from 3 separate experiments. **(D)** Gpc4 protein expression determined by flow cytometry (MFI: median fluorescence intensity), using a Gpc4-directed antibody, on freshly harvested LMC subsets, from either saline-treated (n=6) or IAV-infected (n=7) (300 TCID50) lungs (8 d.p.i). Gating strategy shown in Figure S2 (total lung). Graph contains all values from 2 separate experiments. **(E)** Sdc2 protein expression determined by flow cytometry (MFI: median fluorescence intensity), using a Sdc2-directed antibody, on freshly harvested LMC subsets, from either saline-treated (n=6) or IAV-infected (n=7) (300 TCID50) lungs (8 d.p.i). Gating strategy shown in Figure S2 (total lung). Graph contains all values from 2 separate experiments. Standard error displayed on graphs; n.s: not significant, *: 0.01<p<0.05, **: 0.001<p<0.01, ***: 0.0001<p<0.001, ****: p<0.0001.

In order to experimentally address the importance of these HS core proteins for Areg signaling in LMC, we utilized an siRNA-mediated knockdown approach **(Fig. 4B)**. Treatment of LMC with *Gpc4*-targeting siRNA significantly reduced Areg signaling as compared to LMC treated with control (non-targeting) or *Sdc2*-targeting siRNAs, whereas EGF signaling was unaffected **(Fig. 4C)**.

We then sought to determine the expression levels of glypicans and syndecans on different LMC subsets *in vivo*. We hypothesized that certain cell surface HS core proteins might be induced by specific LMC subsets in the context of tissue damage. Thus, we stained for Sdc2 and Gpc4 in lungs undergoing models of lung damage. Strikingly, we found that Gpc4 is highly upregulated on Col14-LMC in the context of IAV infection **(Fig. 4D)**; no change from baseline was seen in Col13-LMC, while Hhip-LMC show a change of much smaller magnitude. Conversely, Sdc2 did not show similar increases; in fact, it showed significant downregulation on Col14-LMC and Col13-LMC during IAV infection **(Fig. 4E)**. In the context of bleomycin infection, Gpc4 was found to be upregulated on all mesenchymal subpopulations **(Fig. S5C)**, suggesting that different sources of damage may influence Areg responsivity of particular LMC subsets. These results indicate that Gpc4 upregulation in the context of IAV infection is specifically restricted to the Col14-LMC subset.

### HS affects Areg-related tissue repair pathways in vivo

To test the necessity of HS for proper Areg-mediated tissue repair *in vivo*, we conditionally deleted *Ext1* in lung LMC that express *Col1a2* by crossing Col1a2-CreER mice with *Ext1*^fl/fl^ mice [27, 28]. To assess knockout efficiency in various cell types, we sorted lung cells from tamoxifen (TMX)- treated *Ext1*^fl/fl^ control or Col1a2-CreER^+^ *Ext1*^fl/fl^ mice and performed PCR for the targeted genomic region in the *Ext1* gene **(Fig. S6A)**. Among broad cell populations, mesenchymal cells show the highest level of *Ext1* deletion compared to hematopoietic, endothelial, and epithelial cells. Among LMC subpopulations, only Col14-LMC show maximum the level of deletion, in line with our previous studies showing that the *Col1a2* gene is most highly expressed in this subpopulation [3].

To confirm the specificity and efficacy of HS KO we harvested lungs from TMX-administered control and Col1a2-CreER^+^ *Ext1*^fl/fl^ (HS^cKO^) mice and performed flow cytometry with the 10E4 HS-targeted antibody. Only Col14-LMC show a significant reduction in HS staining in HS^cKO^ mice **(Fig. 5A)**. Interestingly, contrary to the accepted narrative in the field of HS study that tissue cells have ubiquitous expression of HS, we found that staining for HS was in fact fairly low in Col14-LMC as compared to other cell types. Strikingly, when we performed HS staining on lungs of mice undergoing the lung damage models described below (IAV and bleomycin), we found Col14-LMC substantially upregulate levels of HS in the setting of tissue damage **(Fig. 5B, S6D)**. Among broader cell types, this upregulation does not occur for hematopoietic cells, endothelial cells, epithelial cells, or even in total mesenchymal cells; among other LMC subsets, Col13-LMC actually show a significant decrease in HS upon lung damage, and while a significant increase is apparent for Hhip-LMC, it is of lesser magnitude/to a lower maximal level than for Col14-LMC. With regards to HS downregulation in HS^cKO^ mice undergoing these models, Col14-LMC show the greatest loss of HS in HS^cKO^ mice compared to control animals. Hematopoietic, endothelial, and epithelial cells show no loss of HS, while significant differences are apparent in total mesenchymal cells. Col13-LMC (during IAV infection only) and Hhip-LMC (during bleomycin treatment only) exhibit significant reductions in surface HS, but to a lesser degree than that seen for Col14-LMC **(Fig. 5B, S6D)**. Thus, we conclude from these data that *(1)* HS is in fact a *damage-inducible* modality for Col14-LMC, rather than a ubiquitously expressed mediator, and *(2)* the Col1a2-CreER driver most effectively targets Col14-LMC among mesenchymal subtypes.

**Figure 5.**
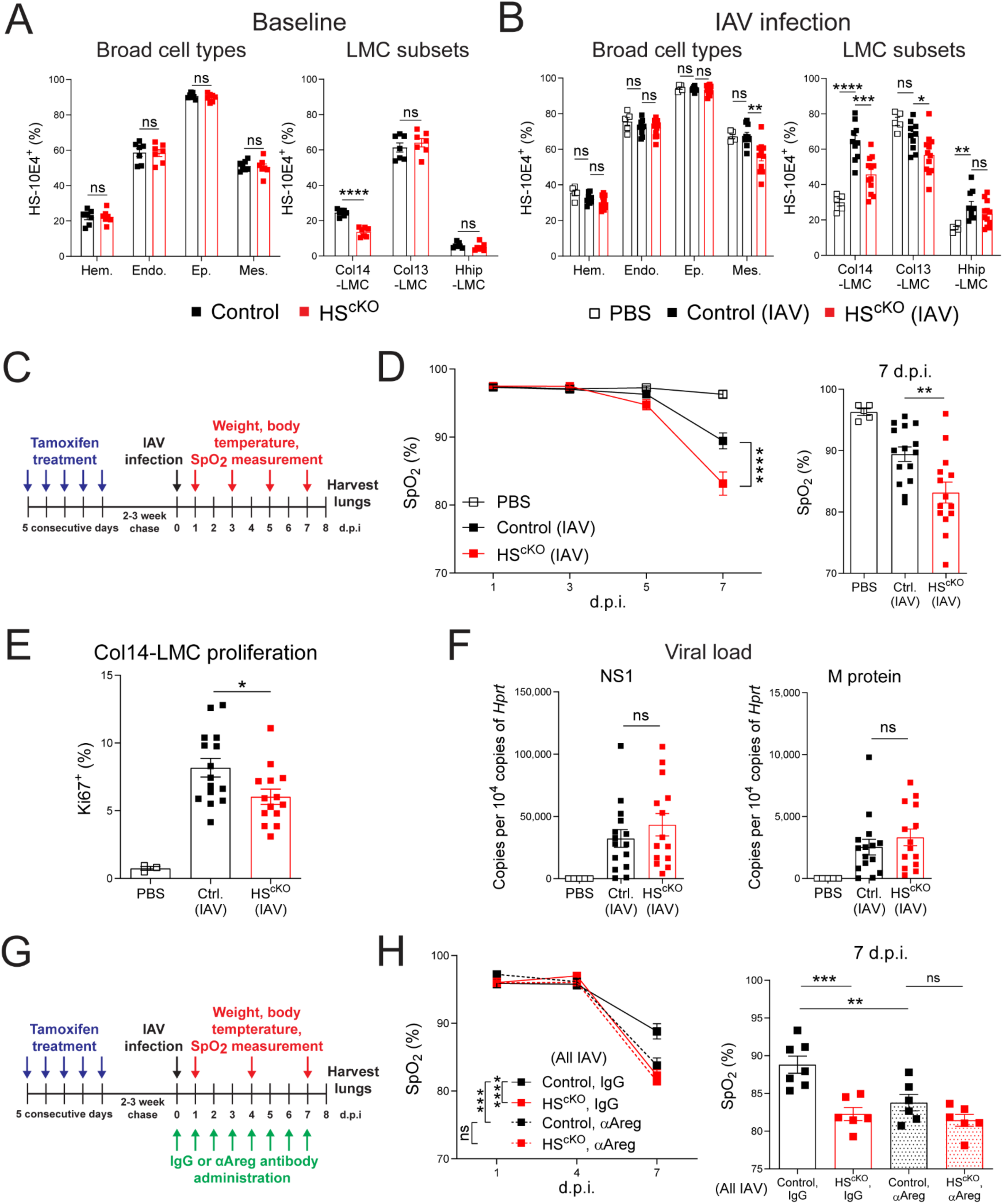
HS affects Areg-related tissue repair pathways in vivo. **(A)** HS presence, assessed by flow cytometric staining for the HS-directed 10E4 antibody, on indicated cell populations from control (n = 7) and HS^cKO^ (n=7) mouse lungs at baseline (all TMX-treated). Gating strategy shown in Figure S2 (total lung). Gating based on FMO controls. Graphs contain all values from 3 separate experiments. **(B)** HS presence, assessed by flow cytometric staining for the HS-directed 10E4 antibody, on indicated cell populations from mock-infection control (PBS) mice (n=5), IAV-infected (100 TCID50) control mice (n=10), and IAV-infected HS^cKO^ mice (n=12) at 8 d.p.i. (all TMX-treated). Gating strategy shown in Figure S2 (total lung). Gating based on FMO control. Graphs contain all values from 3 separate experiments. **(C)** Experimental schematic of IAV model used in (D–F). **(D)** Blood oxygen saturation (SpO_2_) levels in mock-infection control (PBS) mice (n=5), IAV-infected (100 TCID50) control mice (n=15), and IAV-infected HS^cKO^ mice (n=14) (all TMX-treated), with the same mice undergoing repeated measurement at 1, 3, 5, and 7 d.p.i.. All SpO_2_ values from 7 d.p.i. are shown in graph to the right. Graphs contain all values from 3 separate experiments. **(E)** Proliferation induction, assessed via flow cytometry using a Ki67-directed antibody, in Col14-LMC from lungs of mice in (D). Gating strategy for Col14-LMC shown in Figure S2 (total lung). Gating based on FMO control. Graph contains all values from 3 separate experiments. **(F)** qPCR of IAV viral NS1 and M protein in lungs from mice in (D-E). Expression values computed as copies of target gene per 10,000 copies of housekeeping gene (*Hprt*). Graphs contain all values from 3 separate experiments. **(G)** Experimental schematic of IAV model used in (H); daily αAreg or normal goat IgG (control) administration done with 5 μg in 200 ul PBS i.p.. **(H)** SpO_2_ levels in IAV-infected control mice treated with IgG control antibody (n=7) or anti-Areg antibody (n=6) and HS^cKO^ mice treated with IgG control antibody (n=6) or anti-Areg antibody (n=6) (all TMX-treated), with the same mice undergoing repeated measurement at 1, 4, and 7 d.p.i.. All SpO_2_ values from 7 d.p.i. are shown in graph to the right. Graphs contain all values from 3 separate experiments. Standard error displayed on graphs; n.s: not significant, *: 0.01<p<0.05, **: 0.001<p<0.01, ***: 0.0001<p<0.001, ****: p<0.0001.

Given that prior work from our group has shown that Treg cell–derived Areg signaling to Col14-LMC is critical for tissue repair following influenza infection [2, 3], we sought to ascertain the role of HS in this interaction using HS^cKO^ mice **(Fig. 5C)**. Similar to previous studies in Treg cell–specific Areg KO mice [2], we found no changes in weight loss or body temperature between IAV-infected control and HS^cKO^ mice **(Fig. S6B, S6C)**. However, at the level of blood oxygen saturation (SpO_2_), we found a significant decrease over the course of disease – with maximal differences seen at 7 d.p.i. – in IAV-infected HS^cKO^ mice compared to controls **(Fig. 5D)**. We additionally saw a significant decrease in the proliferation of Col14-LMC in IAV-infected HS^cKO^ mice compared to IAV-infected control animals at 8 d.p.i. **(Fig. 5E)**, which is likely indicative of a failure to properly activate these cells in the context of HS deficiency. Importantly, we saw no differences in overall viral load **(Fig. 5F)** or immune cell infiltrate between genotypes (besides slight differences in NK cell and γδ T cells) **(Fig. S7)**. Together, these results indicate that the deficiencies seen here in HS^cKO^ mice are the result of aberrant tissue repair, phenocopying what we previously observed with Treg cell–specific *Areg* deletion or deletion of *Egfr* on Col14-LMC [2, 3].

To investigate whether this failure to properly activate tissue repair activity is indeed Areg dependent, we utilized an antibody blocking approach that has previously shown efficacy in other Areg-related lung repair models [29] **(Fig. 5G)**. As compared to IgG-treated controls, mice treated with αAreg antibody exhibited reduced blood oxygen saturation following IAV infection **(Fig. 5H)**, mirroring the decrease seen in IAV-infected HS^cKO^ mice **(Fig. 5D)**. No further reduction in blood oxygen saturation was observed upon IAV infection of HS^cKO^ mice treated with αAreg antibody. These results imply that the defects in tissue repair seen in HS^cKO^ mice is indeed Areg-dependent.

To extend our investigations using this mouse line to other forms of lung damage, we utilized the bleomycin model of sterile lung damage, which causes extensive alveolar damage and lung fibrosis. Despite the profound loss of HS seen on Col14-LMC in this model **(Fig. S6D)**, disease induction – as indicated by both weight loss **(Fig. S6E)** and staining for α-smooth muscle actin (α-SMA) on LMC as a proxy for fibrosis **(Fig. S6F)** – was not significantly altered.

### HS on Col14-LMCs confers proper responsiveness to Treg cell–derived signals

Finally, to explore the impact of Col14-LMC HS on Treg cell–mediated signaling, we utilized a Col14-LMC/Treg cell co-culture system wherein Col14-LMC from control and HS^cKO^ mice were sorted and co-cultured with primary Treg cells from IAV-infected lungs (8 d.p.i.) **(Fig. 6A)**. Staining for HS in Col14-LMC showed an essentially complete loss of HS on cells from HS^cKO^ mice compared to controls **(Fig. 6B)**. Transcription of *Lif*, previously described to be Treg cell inducible [3], was significantly reduced in HS^cKO^ Col14-LMC in following incubation with Treg cells, compared to control Col14-LMC **(Fig. 6C)**. This reduction was also observed for *Il6*, for which expression has previously been shown to be Treg cell–inducible in astrocytes [30] **(Fig. 6C)**. Following incubation with Treg cells, angiogenesis mediator *Vegfa* showed a trend toward increased expression in control Col14-LMC, while no increase was seen in HS^cKO^ cells expression **(Fig. 6C)**. Notably, for all genes analyzed here, baseline (without incubation with Treg cells) decreases in transcription were observed for HS^cKO^ Col14-LMC vs. control Col14-LMC **(Fig. 6C)**; this may be indicative of a baseline deficiency in the tissue repair capabilities of Col14-LMC that lack HS.

**Figure 6.**
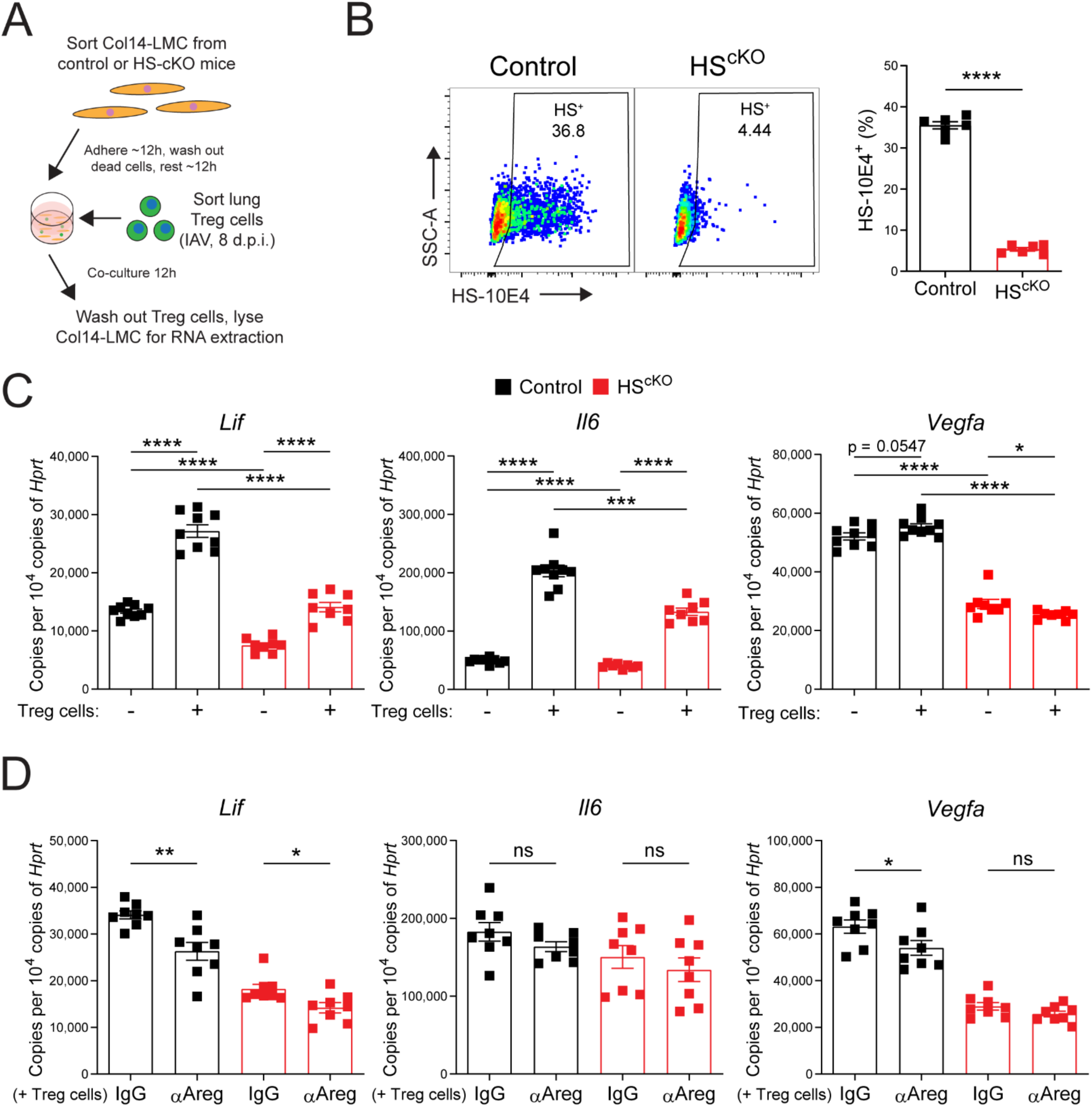
HS on Col14-LMCs confers proper responsiveness to Treg cell–derived signals. **(A)** Experimental schematic for Col14-LMC/IAV-induced lung Treg cell co-culture experiments. **(B)** HS presence, assessed by flow cytometric staining for the HS-directed 10E4 antibody, on Col14-LMC sorted from control (n=6) and HS^cKO^ (n=6) mouse lungs (baseline) and cultured for 48h (non-adherent cells washed out and media replaced at 12h), with the gating/sorting strategy described in Figure S2 (negative bead enriched). Representative flow cytometry plots shown. Gating based on FMO control. Percent staining positive displayed in plots. Graph contains all values from 2 separate experiments. **(C)** qPCR for IAV (300 TCID50) lung Treg cell-inducible genes *Lif*, *Il6*, *Il11*, and *Vegfa* from control (n=9) and HS^cKO^ (n=8) Col14-LMC without inclusion of IAV-induced lung Treg cells, or control (n=9) and HS^cKO^(n=8) Col14-LMC with inclusion of IAV-induced lung Treg cells. Expression values computed as copies of target gene per 10,000 copies of housekeeping gene (*Hprt*). Expression values were normalized across experiments. Graphs contain all values from 3 experiments. **(D)** qPCR for Treg cell-inducible genes *Lif*, *Il6*, *Il11*, and *Vegfa* from control (n=8) and HS^cKO^ (n=8) Col14-LMC co-cultured with IAV-induced lung Treg cells and control IgG antibody (2 μg/ml), or control (n=8) and HS^cKO^ (n=8) Col14-LMC co-cultured with IAV-induced lung Treg cells and αAreg antibody (2 μg/ml). Expression values computed as copies of target gene per 10,000 copies of housekeeping gene (*Hprt*). Expression values were normalized across experiments. Graphs contain all values from 3 separate experiments. Standard error displayed on graphs; n.s: not significant, *: 0.01<p<0.05, **: 0.001<p<0.01, ***: 0.0001<p<0.001, ****: p<0.0001.

To test whether these phenotypes were dependent on Treg cell–derived Areg, αAreg or IgG control antibodies were included in co-cultures. In this setting, we found that *Lif* expression was significantly decreased in the presence of αAreg antibody for both control and HS^cKO^ Col14-LMC, but to a lesser magnitude in the latter **(Fig. 6D)**. Additionally, *Vegfa* gene expression was significantly reduced upon addition of αAreg in control Col14-LMC co-cultures, while in co-cultures with HS^cKO^ Col14-LMC it was unaltered **(Fig. 6D)**. *Il6* did not show alteration in either control or HS^cKO^ Col14-LMC in this context **(Fig. 6D)**. Ultimately, these experiments highlight a reduction in the responsiveness of Col14-LMC to Treg cells (in a partially Areg-dependent manner) and alterations in the baseline transcriptional state of Col14-LMC when these cells lack HS, which likely underlies deficiencies in their reparative capacity.

## Discussion

In this report, we sought to gain a deeper understanding of the manner that Areg, an immune system–derived growth factor involved in tissue repair during damage, interacts with HS. In doing so, we made several discoveries that serve to expand our understanding of the biology of Areg–HS interactions in the context of repair from lung damage.

Previous reports on HSBPs generally regard HS on cognate cells as an obligate member of a signaling complex involving a receptor, an HS-binding ligand, and HS itself. For example, the FGF family is known to require formation of a ternary complex between itself, HS, and cognate FGFRs for signaling [31-33]. In light of this and previous work on the HS-binding capability of Areg [12, 13], we expected that Areg signaling would be absent in cells where HS has been disrupted. However, by testing for downstream MAPK and AKT-mTOR activation in various HS inhibition scenarios, including a completely HS-deficient cell line, we reveal that these pathways maintain activity even in the absence of proper HS engagement, pointing to the existence of HS-dependent and HS-independent signaling modalities for Areg. We used this finding as an opportunity to explore the transcriptional consequences of this previously unappreciated dichotomy with RNA-seq on an Areg-responsive, reparative mesenchymal cell subpopulation from the lung (Col14-LMC). Strikingly, we found that certain Areg-induced transcriptional pathways in these cells are HS-independent (such as expression of tissue factors, growth factors, and angiogenesis mediators), while others are HS-dependent (such as the Hippo pathway, actin mobilization, intermediate filament induction, and gap junction formation). Based on the *in vivo* data we present herein, we believe that the HS-dependent expression modalities we have identified are critical in mediating proper reparative function in Areg target cells. In support of this, activation of the Hippo pathway effector YAP has been shown to promote tissue regeneration in the context of skin wound healing [34]. Areg’s HS binding domain has also been shown to mediate its localization to cell–cell contacts, which could be related to the ability of cells to alter their orientation, polarity, and communication with other cells by engaging these pathways [35]. Furthermore, the biological implications for the existence of HS-independent and -dependent signaling modalities are widespread; the presence or absence of HS at a given signaling interface could plausibly lead to different outcomes for any HSBP.

Another debate in the HS field addressed here involves whether the HS binding ability of a given HSBP is determined by the core protein on which HS is displayed or the sulfation pattern present on HS. In this regard, cell surface HS core proteins are generally thought to be exchangeable, passive displayers of HS, with differences in binding by varying HSBPs conferred instead by the sulfation pattern present on the displayed HS (as determined by the unique set of sulfation enzymes expressed by a given cell type) [9]. In support of sulfation motifs determining binding of a given HSBP, certain HSBPs have been shown to bind HS sequences with distinct sulfation patterns, for instance in the widely studied antithrombin–HS interaction [36]. Thus, we expected that knockout of specific sulfation enzymes would lead to the most significant reduction in Areg signaling potential. Contrary to this paradigm, we found that deletion of a specific GPI-anchored HS core protein (Gpc1) showed the greatest reduction in Areg signaling, while knockout of sulfation enzymes showed little effect. Importantly, knockout of other cell surface HS core proteins did not mediate the same reduction in Areg signaling as Gpc1; furthermore, in Gpc1 knockout cells, a full complement of HS was still displayed on the cell surface, presumably by other core proteins. This result was validated in primary mouse LMC upon siRNA-mediated knockdown of Gpc4, the only glypican expressed by this cell type. “Pre-binding” experiments in which Areg was exposed to heparin of different sulfation states prior to stimulation of cells further implicate Areg as a generalist, sulfation-agnostic HS binder, showing no heightened affinity for specific sulfation sites. These findings suggest that for each HSBP, careful systematic investigation is necessary to determine the nature of HS binding ability.

Interestingly, glypicans (due to their GPI-linked nature) are localized to lipid rafts in cell membranes [37]. A previous study found that this localization has the effect of sequestering FGF2 (an HSBP) away from its non–lipid raft–localized receptor, thus inhibiting its signaling [37]. However, EGFR is known to preferentially localize to lipid rafts [38], which would theoretically confine it and glypicans to a similar location on the cell membrane and potentiate signaling. While plausibly supported by our results, high magnification imaging studies to observe subcellular localization of glypicans and EGFR would be necessary to confirm this interaction. Moreover, *in vivo* models of specific glypican knockouts, which we did not attempt in this work, could be utilized to explore their roles in Areg/EGFR-mediated tissue repair. If further research is done in this regard, it could provide fresh insight into the biology of EGFR signaling, one of the most well-studied pathways in biology.

One additional paradigm related to HS biology addressed in our work is the assumption that tissue parenchymal cells ubiquitously express HS at baseline [11]. While constitutively high expression levels were apparent on the cancer cell lines used in this study, for tissue cells from uninfected mouse lungs, we found that HS levels were in fact quite variable, including relatively low expression on Col14-LMC, a subpopulation known to be critical for tissue repair. Surprisingly, we found that HS is highly upregulated on Col14-LMC (but not altered or only slightly upregulated on other cell types) during lung damage. Furthermore, Gpc4, the HS core protein we identified as preferentially mediating Areg signaling in LMC, similarly shows specific upregulation on Col14-LMC compared to other LMC subsets during IAV infection. The idea of HS as a damage-induced mediator of tissue repair on certain cell subsets runs counter to the concept of “ubiquitous” expression of HS on tissue cell populations, and suggests potential future research directions wherein it may also be found to alter growth factor signaling in disease-specific contexts. The intracellular signaling pathways leading to HS and Gpc4 upregulation merit further investigation; this could plausibly result from inflammatory signaling within the lung environment, sensing of mediators from damaged epithelium, or direct damage to these cells.

In our *in vivo* investigations, we found that mice with an inducible mesenchymal-specific deletion for HS exhibited a reduced ability to recover blood oxygen saturation (SpO_2_) following IAV exposure, which we mechanistically attributed to improper Areg signaling in HS-deleted Col14-LMC. Notably, there were no differences in weight loss, body temperature, immune infiltrate, or viral load (IAV uses sialic acid, not HS for viral entry [39]). The observed decrease in SpO_2_, coupled with the normalcy of these other factors, supports the concept that the former is the result of deficient tissue repair in this context. To the best of our knowledge, this is the first known report of mesenchymal HS serving a critical role in tissue recovery during viral lung infection. Beyond boosting our biological understanding of tissue responses to and recovery from viral infection in the lung, this research suggests several potential avenues of investigation into whether similar dependency on mesenchymal HS is at play for tissue repair in other organs, in the context of other tissue-damaging agents, and/or for other HSBPs.

While the experiments in this report largely speak to basic aspects of HS–immune system interaction, there are several possible future clinical applications for our findings. The discovery that HS is critical for Areg-mediated tissue reparative signaling in the context of lung damage may point to the possibility for treatments that could upregulate or stabilize the presence of HS during tissue injury. However, this becomes complicated in the case of viral infection, where many human viruses rely on HS for entry, and thus HS upregulation could increase viral infection levels. Relatedly, while deploying HS or heparin as decoy viral receptors has been proposed as a way to inhibit viral spreading in patients [40, 41], their delivery could also have the effect of binding up reparative HSBPs such as Areg during tissue recovery; thus, further work is needed to investigate the ideal temporal deployment of these molecules to curtail viral infection while ensuring proper tissue repair.

In conclusion, we have attempted here to provide experimental insight into HS biology as it relates to the immune-derived tissue mediator Areg, which we believe to be an understudied component of immune cell–non-immune cell interaction during tissue repair. Our findings position Areg as an unconventional HSBP that defies many paradigms held in the glycobiology field, while also highlighting that HS along with certain core proteins can serve as damage-inducible tissue modalities in a way that affects tissue repair mechanisms.

## Materials and Methods

### Mice

Wild type (WT) mice (C57BL/6N) were acquired and bred from Jackson Laboratory stocks (Strain #:005304). These mice or lab-bred descendants were utilized for bulk LMC isolation, Col14-LMC isolation/sorting, and *in vivo*/freshly harvested analysis of LMC populations. Col1a2-CreER mice were acquired from Jackson Laboratory (Strain #:029567), and were previously described [27]. *Ext1*^fl/fl^ mice were a generous gift from the laboratory of Dr. Yu Yamaguchi (Sanford Burnham Prebys), and were previously described [28]. Col1a2-CreER mice were crossed to *Ext1*^fl/fl^ mice; these mice were bred with Col1a2-CreER in a hemizygous fashion (Col1a2-CreER^+^ as referred to in this report), with *Ext1*^fl/fl^ as homozygous. These mice showed no overt physical differences, induction of weight loss, or symptoms of distress either at baseline or following tamoxifen treatment. Col1a2-CreER^+^ Ext1^fl/fl^ mice were tested for “leaky” deletion in *Ext1* (i.e., without tamoxifen induction), which was shown not to occur over multiple litters. Mice were screened for the C57BL/6N WT *Nnt* allele (as opposed to the mutant *Nnt*^C57BL/6J^ allele), with all mice used for experiments containing at least one copy of the WT allele. Foxp3^EGFP^ mice were a generous gift from the laboratory of Dr. Alexander Rudensky (Memorial Sloan Kettering), and were previously described [42].

### Cell lines

Lewis Lung Carcinoma (LLC) cells and Phoenix-ECO cells were obtained from the American Type Culture Collection (ATCC). A549 cells were a generous gift from the laboratory of Dr. Richard Vallee (Columbia University). Ba/F3 cells were a generous gift from the laboratory of Dr. Michael Green (UMass Chan Medical School). LLC cells, A549 cells, and Phoenix-ECO cells were cultured in DMEM with 100x penicillin/streptomycin, 100x GlutaMAX (all Gibco), and 10% fetal bovine serum (FBS) (Corning); cells were cultured on tissue culture treated plates (Corning). Ba/F3 cells were cultured in RPMI with 100x penicillin/streptomycin, 100x GlutaMAX (all Gibco), and 10% FBS (Corning); cells were cultured on non-tissue culture treated plates. For maintenance of Ba/F3 cells, rmIL-3 (Biolegend) was added to media (1 ng/ml). Details for transfection/transduction of cell lines with various methods are outlined in the Supplemental Materials and Methods.

### RNA-seq

Col14-LMC were sorted as described, then plated at 50,000 cells/well in 48 well tissue culture plates. After ∼24h in culture, media was aspirated, dead cells were washed away with 1 wash of 1x DMEM, then media was replaced, with either sodium chlorate (3 mg/ml) or vehicle (water) added. After overnight incubation (16-18h), cells were treated with vehicle (PBS + 1% BSA) or rmAreg (200, 500, or 1000 ng/ml) (R&D Systems). After 4h of incubation, media was aspirated and cells were lysed in Trizol Reagent (Thermo). RNA was extracted by the Columbia Molecular Pathology Shared Resource using the miRNeasy Micro Kit (Qiagen), and RINs were found to be >9.6 via Bioanalyzer analysis. We worked with the Columbia Genome Center to perform RNA-seq, using poly-A pulldown to isolate RNA, with subsequent library preparation with a Nextera XT Kit (Illumina) and sequencing with a NovaSeq 6000 sequencer (Illumina) at 40 million reads. Base calling was done with RTA (Illumina) and bclfastq2 (v2.19) was used to convert BCL to fastq files with adapter trimming. Pseudoalignment was done with kallisto (v0.44.0) from transcriptomic data (Ensembl v96, Mouse GRCm38.p6). Subsequent analysis including differential expression was done using the DESeq2 package in R, implemented with iDEP (v1.13) [43]. Pathway analysis was done with g:Profiler [44] to analyze representation within KEGG pathways. Secondary data analysis was done on published RNA-seq datasets (NCBI GEO GSE99714 [5], GSE103548 [26], GSE169127 [3]). Seurat (v3) was used to analyze single cell RNA-seq data [45].

### Mouse tamoxifen treatment, lung damage model induction, and disease assessment

Animal experiments were approved by Columbia University’s Institutional Animal Care and Use Committee (protocol AC-AABT2656). For tamoxifen-based induction of CreER, tamoxifen (Sigma Aldrich) was diluted in corn oil (Sigma Aldrich) at 20 mg/ml, and shaken overnight at 37°C/250 r.p.m. for dissolution in solution, prior to storage at 4°C (<1 month). Mice were treated for 5 consecutive days with 100 mg/kg tamoxifen solution (100-150 µl per mouse); following final tamoxifen treatment, mice were given a 2-3 week chase period prior to experimentation. Influenza A (IAV) virus (PR8/H1N1) was a generous gift from the laboratory of Dr. Donna Farber (Columbia University). For IAV infection, mice were given ketamine/xylazine for anesthesia, then infected intranasally with 100-300 TCID50 of virus diluted in 1x PBS (as determined by the Farber lab using Madin-Darby canine kidney epithelial cell infection assays). Both male and female mice were used for IAV experiments, and mouse lungs were harvested at 8 days post-inoculation (d.p.i.). Bleomycin (Teva) was diluted in sterile 0.9% saline (0.5 U/ml), and 50 µl per mouse was administered oropharyngeally (∼1 U/kg), following a previously described technique [46]. Briefly, mice were given ketamine/xylazine for anesthesia, then placed on an apparatus suspending them at a 60° angle from horizontal by surgical suture string from their teeth. The tongue was removed from the mouth and held with padded forceps, then the bleomycin solution was pipetted into the back of the mouth, followed immediately by plugging the nose with padded forceps to induce oral inhalation of the solution. Male mice were used for bleomycin experiments, and mouse lungs were harvested at 14 d.p.i.. Littermate, age-matched mice were used for all experiments. For IAV and bleomycin experiments, mice were weighed every 1-3 days. For IAV experiments, mouse body temperature was assessed with a rectal thermometer. For IAV experiments, mouse blood oxygen saturation (SpO_2_) was assessed using a MouseOx Plus Pulse Oximeter (Starr Life Sciences). The area around mouse the mouse neck was shaved and Nair Hair Remover Product was applied to chemically remove residual hair at the time of IAV infection. A Small Mouse Collar Sensor (Starr Life Sciences) was used to assess SpO_2_ on unanesthetized at indicated timepoints during IAV infection progression, with mice placed in a 1 L beaker to restrict movement; assessment was taken for 2-5 min. per mouse at each timepoint, with only high-quality, error-free readings taken and averaged to determine a composite SpO_2_ value. For IAV experiments, mice were disincluded from final analysis if they experienced loss of SpO_2_ to a level <85% at 3-4 d.p.i. (as this is an indication of lung damage from technical issues with administration, not from IAV infection), if they did not experience a loss of body temperature <36.5°C at 7 d.p.i. (as this is evidence of administration issues leading to an unproductive infection), or if there was visible failure to uptake IAV intranasally during infection; 6 of 80 total IAV-infected mice in experiments were excluded via these criteria (not including mice for bulk sorting of IAV-induced Treg cells). At endpoint of disease models, lung tissue was perfused and (for certain experiments) bronchoalveolar lavage (BAL) was performed as described in “Lung processing”. For IAV experiments, middle and inferior lobes were flash frozen for RNA extraction to assess viral infection levels, and superior and post-caval lobes were processed for flow cytometry. Lobes for RNA extraction were added to Trizol (Thermo), a ¼" Ceramic Sphere (MP Biomedicals) was added to tubes, and tissue was homogenized in a FastPrep-24 (MP Biomedicals) for lysis, followed by RNA analysis (see “RNA extraction and qPCR”). For bleomycin experiments, full lungs were processed for flow cytometry.

### Western blotting, RNA extraction/qPCR, lung processing, negative bead enrichment/sorting, flow cytometry, siRNA treatment, Col14-LMC/Treg cell co-culture

Details for these methods are outlined in the Supplemental Materials and Methods. For qPCR analysis in certain experiments, results were calculated as copies of the gene of interest per copies of housekeeping gene (*Hprt*) using a previously described method [47].

### Statistics and data analysis

GraphPad Prism (v10.1.2) was used for all statistical analyses and graphing. For western blots, flow cytometry, SpO_2_, or qPCR analysis where two groups were compared, two-tailed unpaired Student’s t-tests were used. For western blot analysis where three or more groups were compared, one-way ANOVA was used. For post-hoc RNA-seq analysis of individual genes where three or more groups were compared, the Kruskal-Wallis test was used. For longitudinal weight loss, body temperature, or SpO_2_ analysis, two-way repeated measure ANOVA was used. Statistical significance was determined at p<0.05, with further levels of significance reported in figure legends. Sample size estimation was determined based on previous studies. FlowJo (v10), Microsoft Excel, Microsoft Powerpoint, SnapGene Viewer, Adobe Illustrator, ImageJ, MacVector, and R were used to set up experiments, analyze data, and prepare data.

## Data, Materials, and Software Availability

RNA-seq data associated with this manuscript have been deposited in NCBI’s Gene Expression Omnibus under GEO accession number GSE263616. We report no original source code from this manuscript. All other data have been included in the manuscript or supplemental information.

## Acknowledgements

We thank several Columbia University Core Facilities for their contributions to this work: the Microbiology & Immunology Shared Resources, Columbia Stem Cell Initiative (CSCI) Flow Cytometry, the Columbia Genome Center, and the Herbert Irving Cancer Comprehensive Cancer Center Molecular Pathology Shared Resource (MPSR), funded in part through the NIH/NCI Cancer Center Support Grant P30CA013696. We thank Dr. Alexander Rudensky for contribution of Foxp3^EGFP^ mice. We thank Dr. Yu Yamaguchi for contribution of *Ext1*^fl/fl^ mice. We thank Dr. Donna Farber for contribution of influenza A virus (PR8/H1N1). We thank Dr. Richard Vallee for contribution of A549 cells. We thank Dr. Michael Green for contribution of Ba/F3 cells. We thank all other Arpaia lab members, as well as S. Iketani and M. Edwards, for their feedback and comments on this work. This work was supported by NIH grants R21AI149657 and R01HL148718.

## Author contributions

L.F.L. and N.A. designed research; L.F.L., A.S., O.R.R., F.L., and K.d.l.S.-A. performed research; L.F.L. and A.S. analyzed data; L.F.L. and N.A. wrote the paper.

---

**Figure S1.**
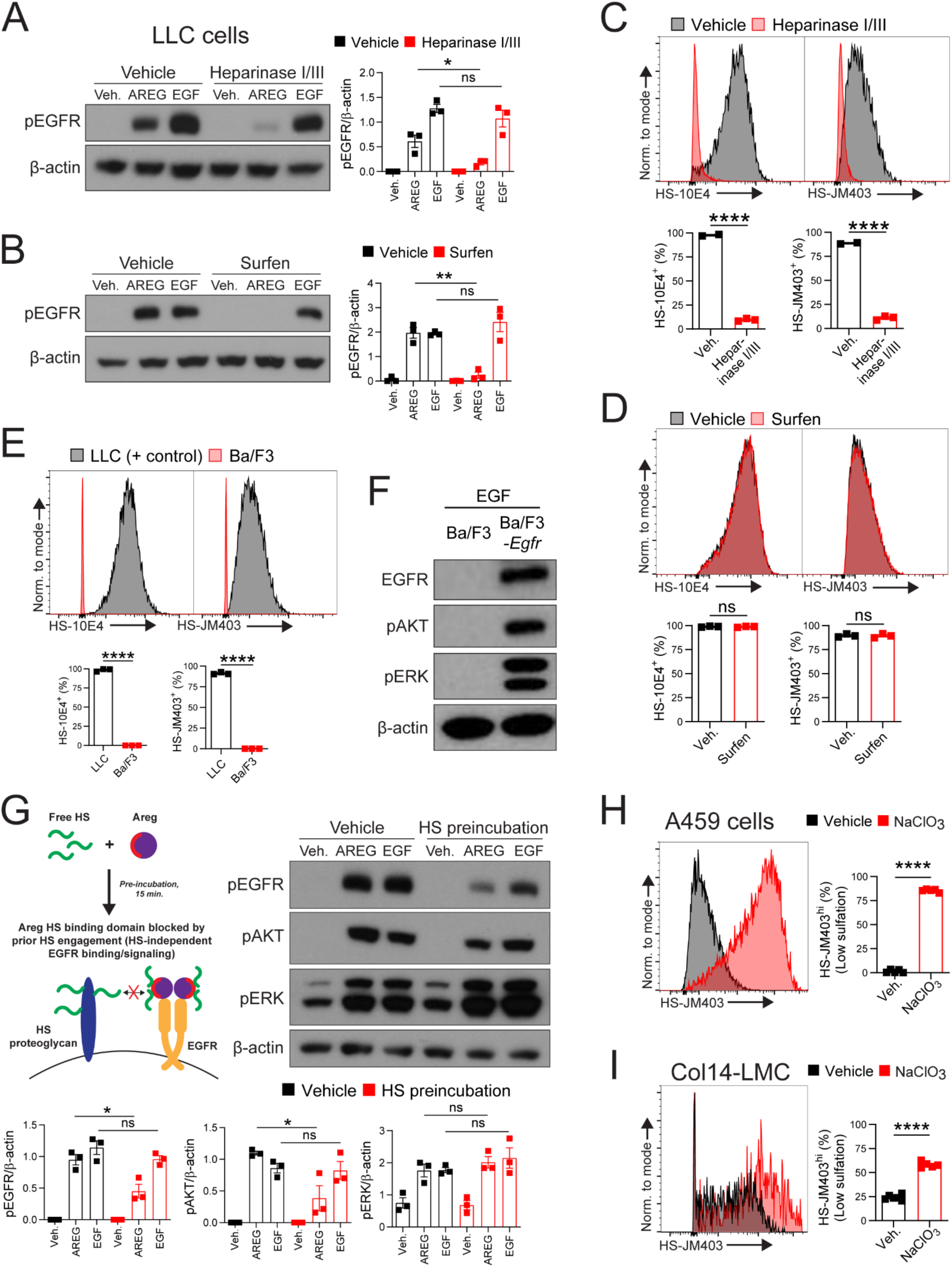
Supplementary data for Figure 1. **(A)** Western blotting for phospho-EGFR (Y1068) and β-actin of vehicle or heparinase I/III-treated (1h) LLC cells, stimulated for 15 min. with vehicle, murine AREG (500 ng/ml), or murine EGF (100 ng/ml). Representative western blots shown. n=3 per condition, graph contains all values from 3 separate experiments. **(B)** Western blotting for phospho-EGFR (Y1068) and β-actin of vehicle or surfen-treated (15 min.) LLC cells, stimulated for 15 min. with vehicle, murine AREG (500 ng/ml), or murine EGF (100 ng/ml). Representative western blots shown. n=3 per condition, graph contains all values from 3 separate experiments. **(C)** Flow cytometry using HS-directed antibodies 10E4 and JM403 on LLC cells treated with vehicle or heparinase I/III (1h). Representative flow cytometry plots shown. Gating based on FMO controls. n=2-3 per condition, graphs contain all values from 2 separate experiments. **(D)** Flow cytometry using HS-directed antibodies 10E4 or JM403 on LLC cells treated with vehicle or surfen (15 min.). Representative flow cytometry plots shown. Gating based on FMO controls. n=3 per condition, graphs contain all values from 2 separate experiments. **(E)** Flow cytometry using HS-directed antibodies 10E4 or JM403 on untreated Ba/F3 cells, with untreated LLC cells used as a positive staining control. Representative flow cytometry plots shown. Gating based on FMO controls. n=3 per cell type, graphs contain all values from 2 separate experiments. **(F)** Western blot showing transfection of Ba/F3 cells with murine *Egfr* (EGFR blot), and EGF (50 ng/ml)-inducible downstream signaling via phospho-AKT (S473) and phospho-ERK (T202/Y204) (β-actin also included). Representative western blots shown from 4 separate experiments. **(G)** Western blotting for phospho-EGFR (Y1068), phospho-AKT (S473), phospho-ERK (T202/Y204), and β-actin of vehicle- or HS-pretreated (15 min.) vehicle, murine AREG (500 ng/ml), or murine EGF (100 ng/ml), subsequently applied to LLC cells (15 min.). Representative western blots shown. n=3 per condition, graphs contain all values from 3 separate experiments. Diagram included of experimental setup (top left). **(H)** Flow cytometry using HS-directed antibody JM403 on A549 cells treated with vehicle or NaClO_3_ (16-18h). Representative flow cytometry plot shown. Gating based on FMO control. n=4 per condition, graph contains all values from 2 separate experiments. I) Flow cytometry using HS-directed antibody JM403 on Col14-LMC treated with vehicle or NaClO_3_ (16-18h). Representative flow cytometry plot shown. Gating based on FMO control. n=5 per condition, graph contains all values from 2 separate experiments. Standard error displayed on graphs; n.s: not significant, *: 0.01<p<0.05, **: 0.001<p<0.01, ***: 0.0001<p<0.001, ****: p<0.0001.

**Figure S2.**
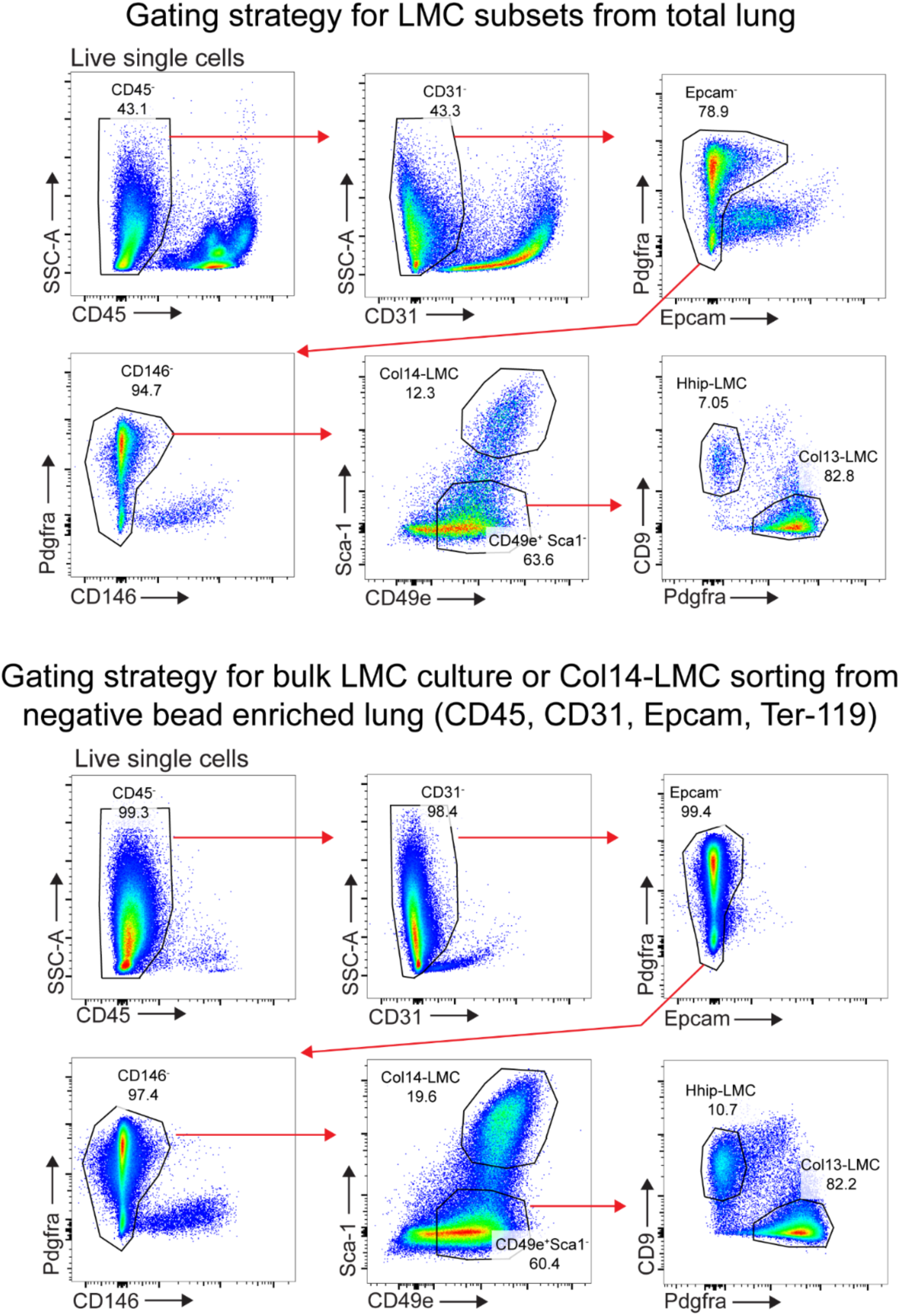
Flow cytometry gating strategies for mouse LMC. Flow cytometry gating strategy for staining of LMC subsets from baseline WT mouse lung, and from negative bead enriched baseline WT mouse lung. Negative bead enrichment was done with biotinylated antibodies towards Ter-119, CD45, CD31, and Epcam, to remove red blood cells, hematopoietic cells, endothelial cells, and epithelial cells, respectively. Flow cytometry analyses of different LMC populations were done using the panel from total lung (no enrichment), while bulk LMC (cultured directly or cell sorting of Col14-LMC) for experiments was done from the negative bead-enriched lung panel. Percent of previously gated population displayed in plots.

**Figure S3.**
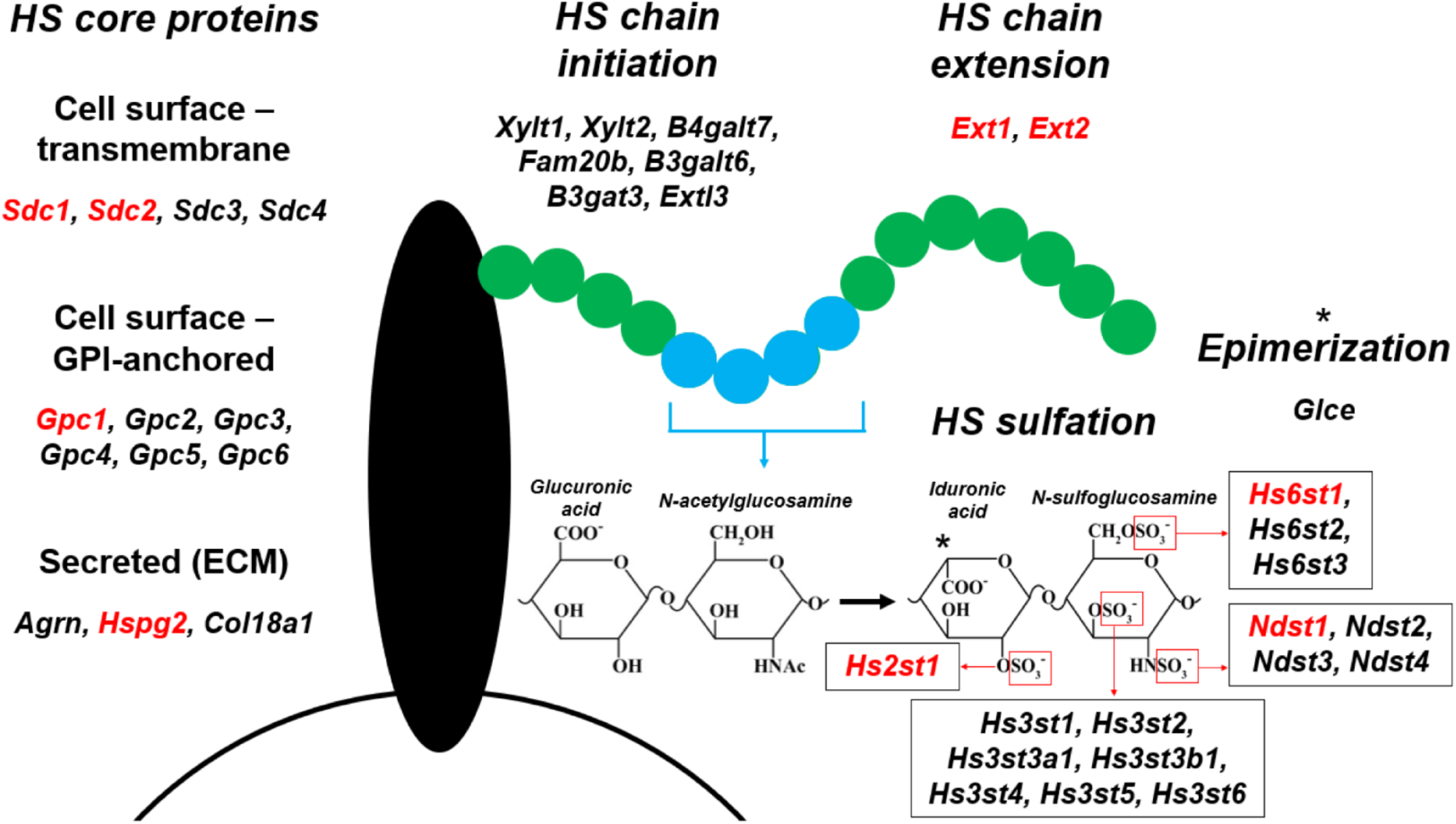
Summary of HS-related genes and their functions. Schematic of HS-synthesis related genes, with those included in LLC cell KO panel in red. HS must be attached to “core proteins” at specific serine sites along their protein structure – either cell surface proteins – syndecans (transmembrane) or glypicans (GPI-anchored) – or secreted ECM proteins. While these core proteins are trafficked through the Golgi apparatus, a series of enzymes catalyzes the addition of the specific sugar moiety combination that initiates all HS chains. Then, the critical enzymes Ext1 and Ext2 are nonredundantly responsible for adding the repeated sugar subunit of glucuronic acid and N-acetylglucosamine to the HS chain. The HS chain upon formation lacks sulfation; a series of enzymes, also operating in the Golgi apparatus, operate on disparate regions of the HS chain to add sulfation groups, the negative charge of which largely confer signaling alteration potential to HS towards positively charge HS binding protein domains. These enzymes are thought to operate in sequential fashion (i.e., sulfation must occur at one site before it can occur at the next), in this order: the N-sulfotransferase enzyme group (Ndst1-4) exchanges an acetyl group for a sulfate group in N-acetylglucosamine (creating N-sulfoglucosamine); Glce epimerizes the indicated carbon in glucuronic acid (creating iduronic acid); the 2-O-sulfotransferase enzyme (Hs2st1) adds a sulfate group at the indicated hydroxyl group in iduronic acid; the 6-O-sulfotransferase enzyme group (Hs6st1-3) adds a sulfate group at the indicated hydroxyl group in N-sulfoglucosamine; and the 3-O-sulfotransferase enzyme group (Hs3st1-6) adds a sulfate group at the indicated hydroxyl group in iduronic acid. HS contains significant heterogeneity in their sulfation profile along each individual chain, with alternating regions of little or no sulfation and regions with high sulfation.

**Figure S4.**
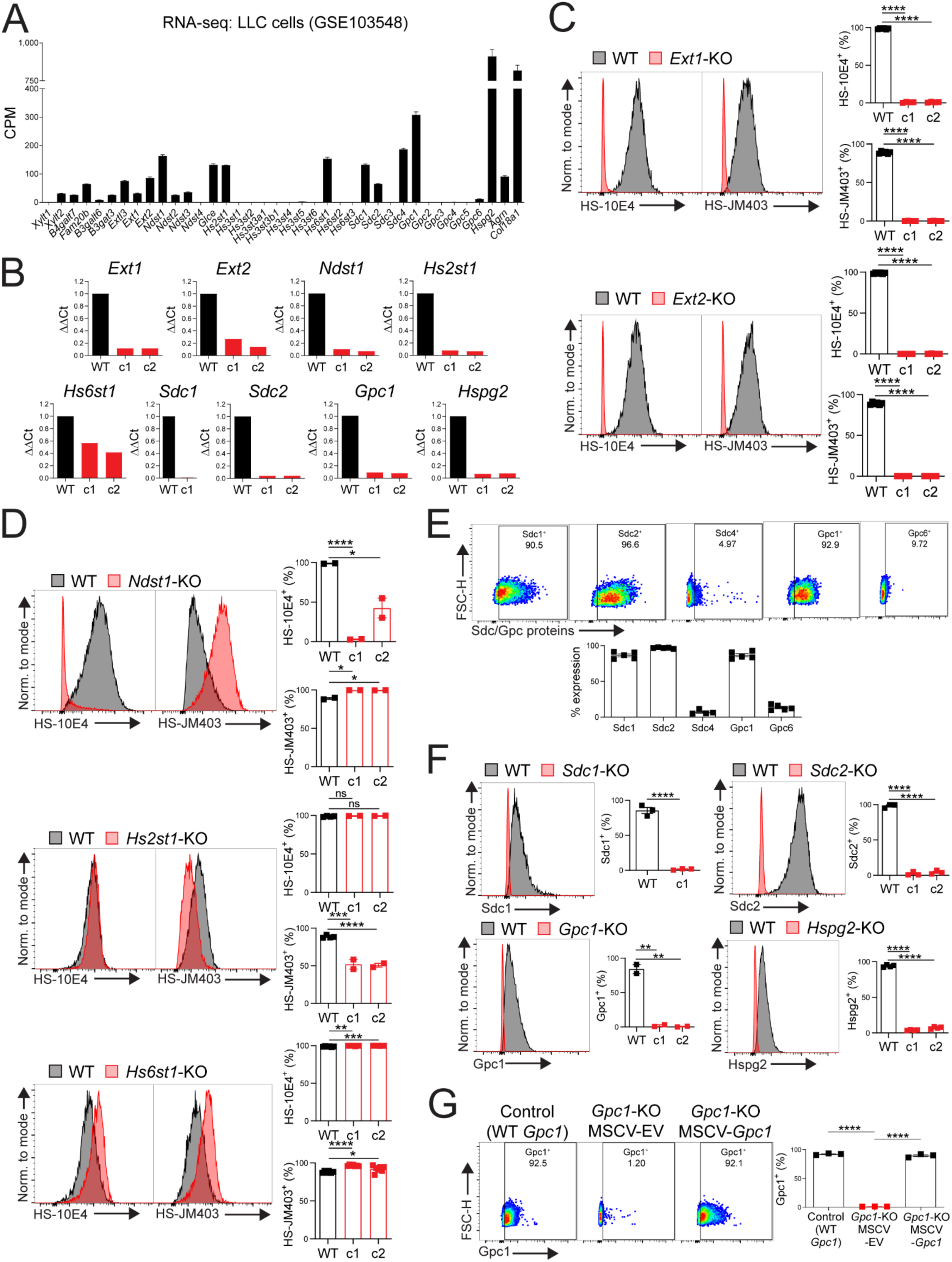
Supplementary data for Figure 3. **(A)** Data from publicly available bulk RNA-seq of LLC cells (GSE103548), of HS synthesis-related genes. CPM: Counts per million. **(B)** qPCR on WT LLC cells or LLC cell KO sublines for genes targeted by CRISPR-Cas9 in LLC cell KO panel for HS-related genes. Expression values were computed as ΔΔCT values, compared to WT. c1/c2: clone 1/clone 2 (i.e., different single cell subcloned lines). Graphs contain all values from 1 experiment per subline. **(C)** Flow cytometry using HS-directed antibodies 10E4 and JM403 on WT vs. *Ext1*- or *Ext2*-KO LLC cell sublines. Representative flow cytometry plots shown. Gating based on FMO controls. n=4-6 per clonal subline, graphs contain all values from 3 separate experiments. **(D)** Flow cytometry using HS-directed antibodies 10E4 and JM403 on WT vs. *Ndst1*-KO, *Hs2st1*-KO, or *Hs6st1*-KO LLC cell sublines. Representative flow cytometry plots shown. Gating based on FMO controls. n=2-7 per clonal subline, graphs contain all values from 2-4 separate experiments. **(E)** Flow cytometry using antibodies targeting Sdc1, Sdc2, Sdc4, Gpc1, or Gpc6 on WT LLC cells. Representative flow cytometry plots shown. Gating based on FMO controls. Percent staining positive displayed in plots. n=4-5 per target, graph contains all values from 2-4 separate experiments. **(F)** Flow cytometry using antibodies targeting Sdc1, Sdc2, or Gpc1 on WT vs. *Sdc1*-, *Sdc2*-, *Gpc1*-KO, or *Hspg2*- KO LLC cell sublines. Representative flow cytometry plots shown. Gating based on FMO controls. n=2-4 per clonal subline, graphs contain all values from 2-3 separate experiments. **(G)** Flow cytometry using an antibody targeting Gpc1 on CRISPR-Cas9 control (WT *Gpc1*), *Gpc1*-KO LLC cells transduced with empty vector (EV) MSCV retrovirus (MSCV-EV), or *Gpc1*-KO LLC cells transduced with MSCV retrovirus with *Gpc1* mRNA (MSCV-*Gpc1*). Representative flow cytometry plots shown. Gating based on FMO control. Percent staining positive displayed in plots. n=3 per group, graph contains all values from 3 separate experiments. Standard error displayed on graphs; n.s: not significant, *: 0.01<p<0.05, **: 0.001<p<0.01, ***: 0.0001<p<0.001, ****: p<0.0001.

**Figure S5.**
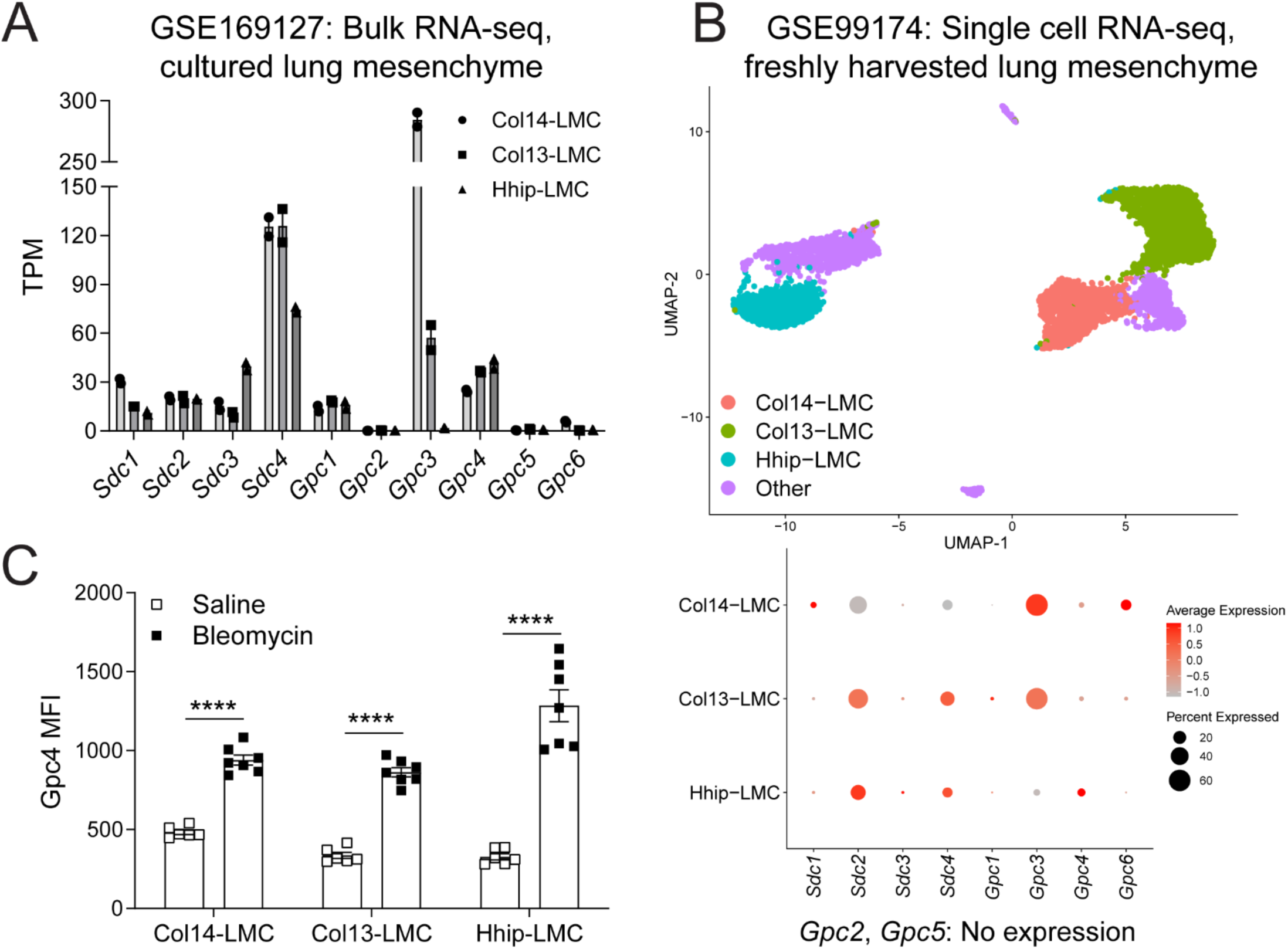
Supplementary data for Figure 4. **(A)** Data from publicly available bulk RNA-seq of lung mesenchyme subsets (GSE169127), for HS core protein genes. **(B)** Data from publicly available scRNA-seq of lung mesenchyme (GSE99714), for HS core protein genes. **(C)** Gpc4 protein expression determined by flow cytometry (MFI: median fluorescence intensity), using a Gpc4-directed antibody, on freshly harvested LMC subsets, from either saline-treated (n=6) or bleomycin-treated (n=7) (1 U/kg) lungs (14 d.p.i). Gating strategy shown in Figure S2 (total lung). Graph contains all values from 2 separate experiments. Standard error displayed on graphs; n.s: not significant, *: 0.01<p<0.05, **: 0.001<p<0.01, ***: 0.0001<p<0.001, ****: p<0.0001.

**Figure S6.**
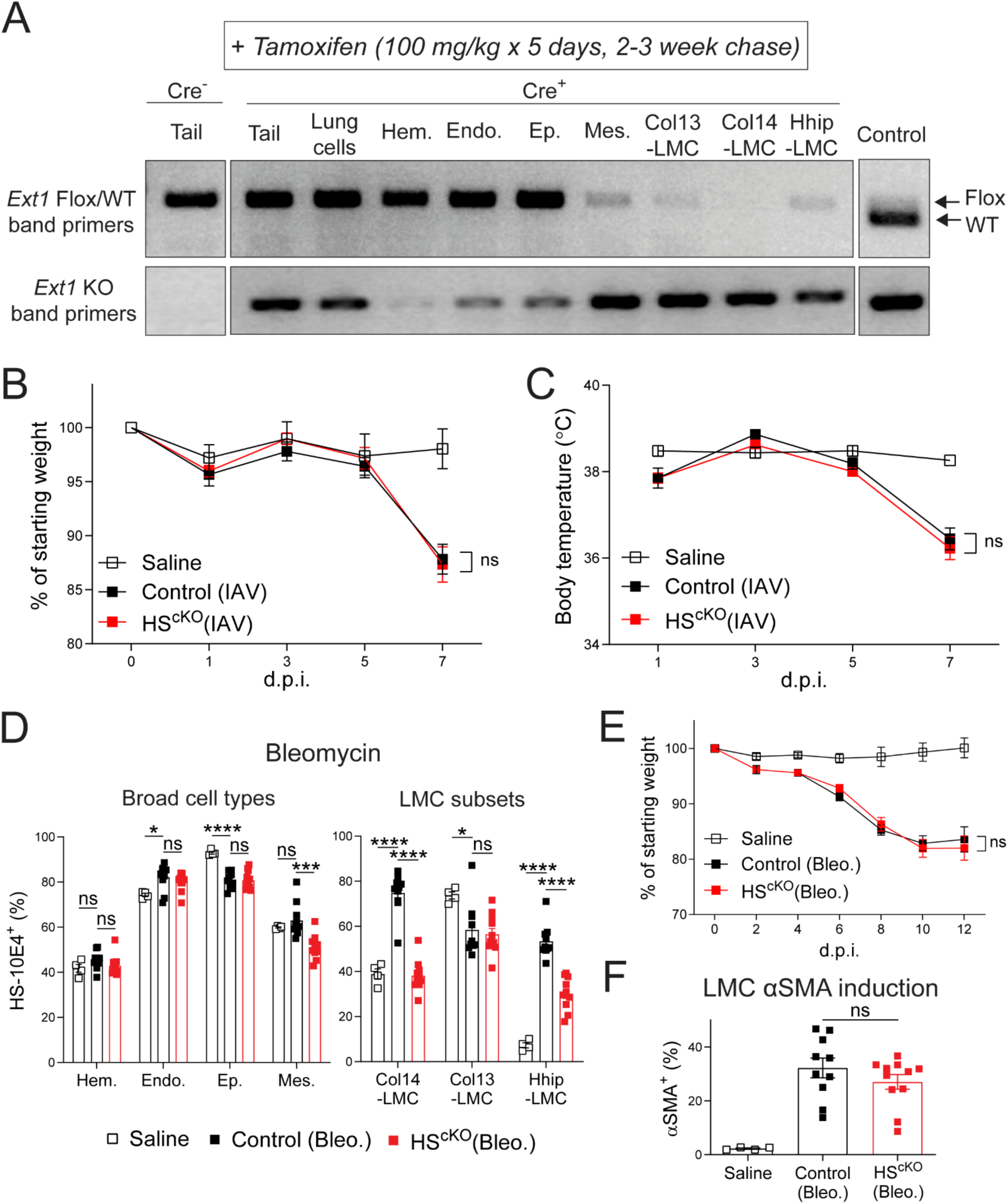
Supplementary data for Figure 5. **(A)** PCR-based genotyping of DNA from tails, bulk lung cells, and flow cytometry-sorted cell populations from TMX-administered control and HS^cKO^ mice at baseline. Top gel: PCR with primers targeting WT mouse *Ext1* (WT band), or *Ext1* with LoxP sites added (Flox band). Bottom gel: PCR with primers targeting LoxP recombination-mediated, exon 1-excised *Ext1* (KO band). Representative gel shown from 2 separate experiments. **(B)** Percent starting weight for mock-infection control (PBS) mice (n=5), IAV-infected (100 TCID50) control mice (n=15), and IAV-infected HS^cKO^ mice (n=14) (all TMX-treated) at 0, 1, 3, 5, and 7 d.p.i.. Graph contains all values from 3 separate experiments. **(C)** Body temperature of mock-infection control (PBS) mice (n=5), IAV-infected (100 TCID50) control mice (n=15), and IAV-infected HS^cKO^ mice (n=14) (all TMX-treated) at 1, 3, 5, and 7 d.p.i.. Graph contains all values from 3 separate experiments. **(D)** HS presence, assessed by flow cytometric staining for the HS-directed 10E4 antibody, on indicated cell populations from mock-treated control (saline) mice (n=4), bleomycin-treated (1U/kg) control mice (n=11), and bleomycin-treated HS^cKO^ mice (n=12) (all TMX-treated) at 14 d.p.i.. Gating strategy shown in Figure S2 (total lung). Gating based on FMO control. Graphs contain all values from 3 separate experiments. **(E)** Percent starting weight for mock-treated control (saline) mice (n=4), bleomycin-treated (1U/kg) control mice (n=11), and bleomycin-treated HS^cKO^ mice (n=12) (all TMX-treated) at 0, 2, 4, 6, 8, 10, and 12 d.p.i.. Graph contains all values from 3 separate experiments. **(F)** Fibrosis induction, assessed via flow cytometry using a α-smooth muscle actin (αSMA)-directed antibody, in Pdgfra^+^ LMC from mice in (E), at 14 d.p.i.. Gating strategy shown in Fig. S2 (total lung). Gating based on FMO control. Graph contains all values from 3 separate experiments. Standard error displayed on graphs; n.s: not significant, *: 0.01<p<0.05, **: 0.001<p<0.01, ***: 0.0001<p<0.001, ****: p<0.0001.

**Figure S7.**
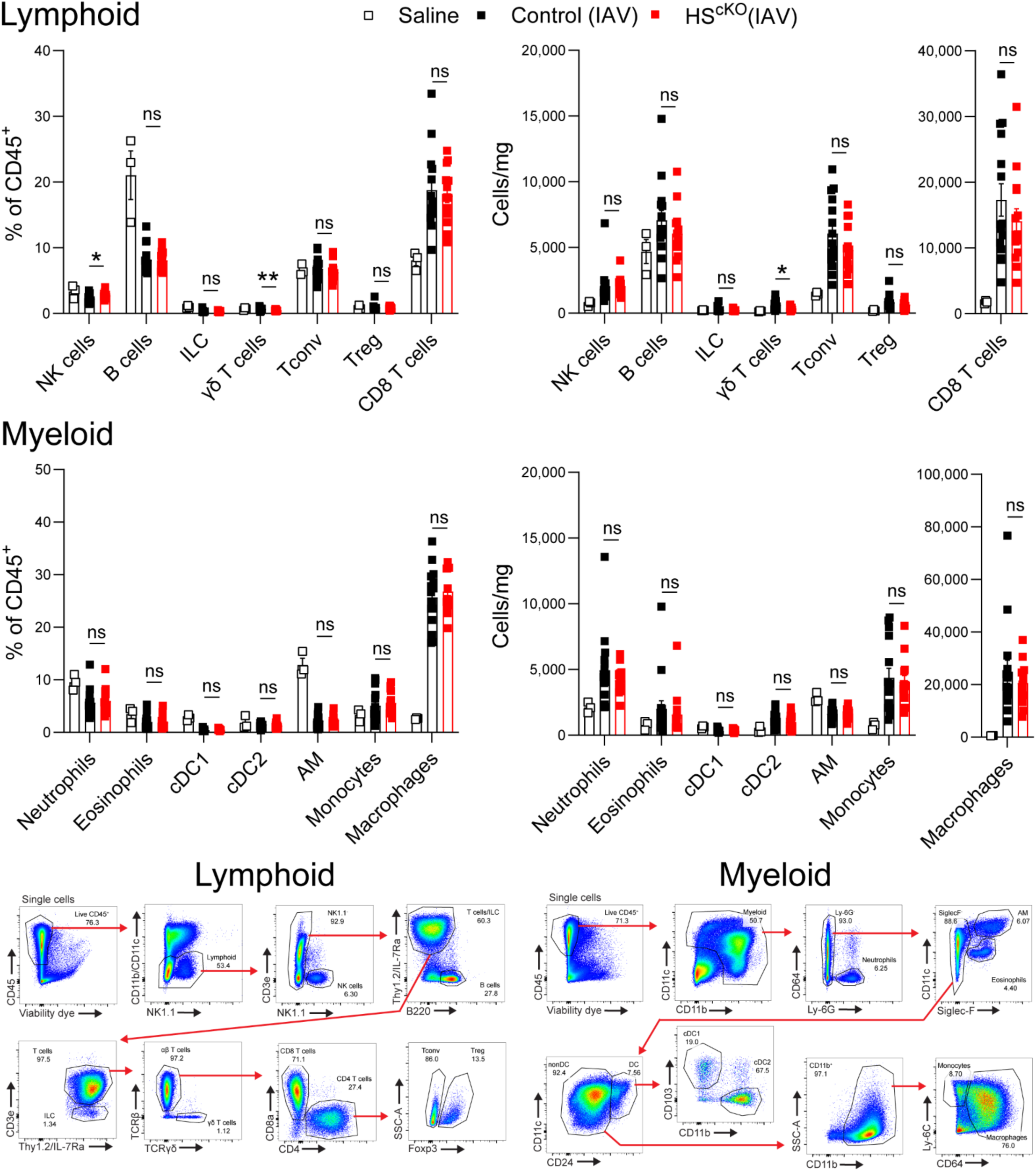
Immune infiltrate in IAV-infected control and HS^cKO^ mouse lungs (Supplementary data for Figure 5) Percentage of all CD45^+^ cells and cell counts per lung weight (determined by flow cytometry) for predominant myeloid and lymphoid immune cell types from mock-infection control (PBS) mice (n=3), IAV-infected (100 TCID50) control mice (n=15), and IAV-infected HS^cKO^ mice (n=14) (all TMX-treated) at 8 d.p.i.. Gating strategies for immune cell type identification shown (below); example plots are from IAV-infected mouse lungs (8 d.p.i.). Percent of previously gated population displayed in plots. Graphs contain all values from 3 separate experiments. Standard error displayed on graphs; n.s: not significant, *: 0.01<p<0.05, **: 0.001<p<0.01, ***: 0.0001<p<0.001, ****: p<0.0001.

## Supplemental Materials and Methods

### Western blotting

For LLC cells and A549 cells, cells were plated at 250,000 cells per well in 6 well tissue culture treated plates, allowed to adhere for 24h, then washed and kept in starving media (all media components besides FBS) overnight (16-18h) before ligand treatment (15 min.) and lysis. HS-modifying agents used in certain experiments were sodium chlorate (3 mg/ml; added at time of starving media replacement, 16-18h prior to ligand treatment; Sigma-Aldrich), heparinase I/III (2-5 U/ml; added 1h prior to ligand treatment; Sigma-Aldrich), and surfen (5 µg/ml; added 15 min. prior to ligand treatment; EMD Millipore/Calbiochem). For pre-treatment experiments, prior to treatment of cells, ligands were pre-incubated with heparan sulfate (1 µg/ml; Sigma), heparin (1 µg/ml; Sigma), or desulfated heparin variants (2-O-desulfated heparin, 6-O-desulfated heparin, N-desulfated heparin, N-desulfated-reacetylated heparin [1 µg/ml; AMSBio]) for 15 min., then this mixture was applied to target cells. For Ba/F3 cells, cells were washed twice in starving media (all media components besides FBS; no rmIL-3), then plated at 2 million cells per well in 24 well non-tissue culture treated plates and cultured overnight (16-18h) before ligand treatment and lysis. For isolated bulk LMC or Col14-LMC (see “Bulk LMC or CD4 T cell negative bead enrichment, and Col14-LMC and Treg sorting”), cells were plated in mesenchymal cell media, allowed to adhere for 12-24h, then washed and kept in fresh mesenchymal cell media overnight (16-18h) or for 48h (for siRNA experiments) before ligand treatment and lysis. The next morning (LLC, Ba/F3, A549, Col14-LMC) or 48h later (bulk LMC treated with siRNA), cells were treated with vehicle, recombinant mouse Areg (R+D), human AREG (R+D), mouse EGF (Biolegend), or human EGF (R+D), for 15 min., then lysed for protein extraction. For LLC, A549, and Col14-LMC, cells were placed on ice, washed twice with cold 1x PBS to remove dead cells, then lysed by addition of TNT Lysis Buffer (20 mM Tris-HCl [Sigma-Aldrich], 150 mM sodium chloride [Fisher], 1 mM ethylenediaminetetraacetic acid [EDTA] disodium salt dihydrate [Fisher], 1 mM ethyleneglycol bis(2-aminoethyl ether)-N,N,N’,N’ tetraacetic acid [EGTA] [Gold Biotechnology], 1% Triton X-100 [Sigma-Aldrich], pH 8.0) with freshly added Halt Protease and Phosphatase Inhibitor Cocktail (Pierce) and scraping of wells. For Ba/F3 cells (non-adherent), cells were harvested from wells, brought to 10 ml with cold 1x PBS, centrifuged at 550 x g, 4°C, 2 min., supernatant was aspirated, cells were washed again in 10 ml cold 1x PBS, centrifuged/supernatant aspirated again, then lysed in TNT Lysis Buffer with freshly added inhibitors. Lysates were shaken for 30 min. on ice, then centrifuged for 15 min. at maximum speed; the supernatant was then removed and used for western blotting. Lysate supernatants were mixed with 6x SDS Sample Buffer (0.375 M Tris-HCl pH 6.8, 12% sodium dodecyl sulfate [SDS] [Millipore], 60% glycerol [Fisher], 0.6 M dithiothreitol [DTT] [Fisher], 0.06% bromophenol blue [Amresco]), incubated at 95°C for 5 min., cooled to room temperature, and loaded into Bolt 4–12% Bis-Tris Plus Gels (Thermo). SDS-PAGE (in Bolt MOPS SDS Running Buffer [Thermo]) and protein transfer (to Immobilon-P PVDF [Millipore] in Bolt Transfer Buffer [Thermo]) were done using the Mini Blot Module (Thermo). Membranes were washed with 1x TBS, blocked using 5% nonfat dry milk in TBST (1x TBS with 0.1% Tween-20 [Fisher]), washed 3 times with TBST, then stained with primary antibodies in 5% milk or BSA in TBST, overnight at 4°C (β-Actin [clone 8H10D10; Cell Signaling], Phospho-EGF Receptor Tyr1068 [clone D7A5; Cell Signaling], Phospho-Akt Receptor Ser473 [clone D9E; Cell Signaling], Phospho-p44/42 MAPK [Erk1/2] Thr202/Tyr204 [clone D13.14.4E; Cell Signaling], EGF Receptor [clone C74B9; Cell Signaling]). After overnight incubation, membranes were washed 3 times with TBST, then stained with secondary antibodies in 5% milk in TBST for 1h at room temperature (anti-mouse IgG HRP-linked antibody [Cell Signaling], anti-rabbit IgG HRP-linked antibody [Cell Signaling]). Afterwards, membranes were washed 3 times with TBST, then incubated with chemiluminescent substrate (Immobilon Classico Western HRP Substrate [Millipore], Immobilon Forte Western HRP Substrate [Millipore], SuperSignal West Femto Chemiluminescent Substrate [Thermo]). Membranes were then exposed to film, with films developed on a Kodak X-OMAT. Within each experiment, blots for phosphorylated proteins were performed first, with subsequent stripping with Restore Western Blot Stripping Buffer (Thermo) and reprobing for β-Actin as a loading control. Western blots were quantified using ImageJ.

### RNA extraction and qPCR

RNA extraction was done using Trizol Reagent (Thermo) for lysis, followed by chloroform-based separation and precipitation of RNA with isopropanol. RNA amounts were subsequently quantified using Nanodrop (Thermo), and cDNA was created using a qScript cDNA Synthesis Kit (Quanta). qPCR was performed on cDNA using SYBR Green qPCR Master Mix (Thermo) and a Bio-Rad CFX384 qPCR system. Primers were created for this study using Integrated DNA Technology’s PrimerQuest platform, and University of California Santa Clara’s In-Silico PCR platform; all primers were ordered from Integrated DNA Technology and are listed in Table 1. Analysis was done by calculating ΔΔCt values relative to housekeeping gene (*Hprt*), or by calculating the copies of an analyzed gene per copies of housekeeping gene (*Hprt*) using a previously described method (Pfaffl 2001).

**Table 1.**
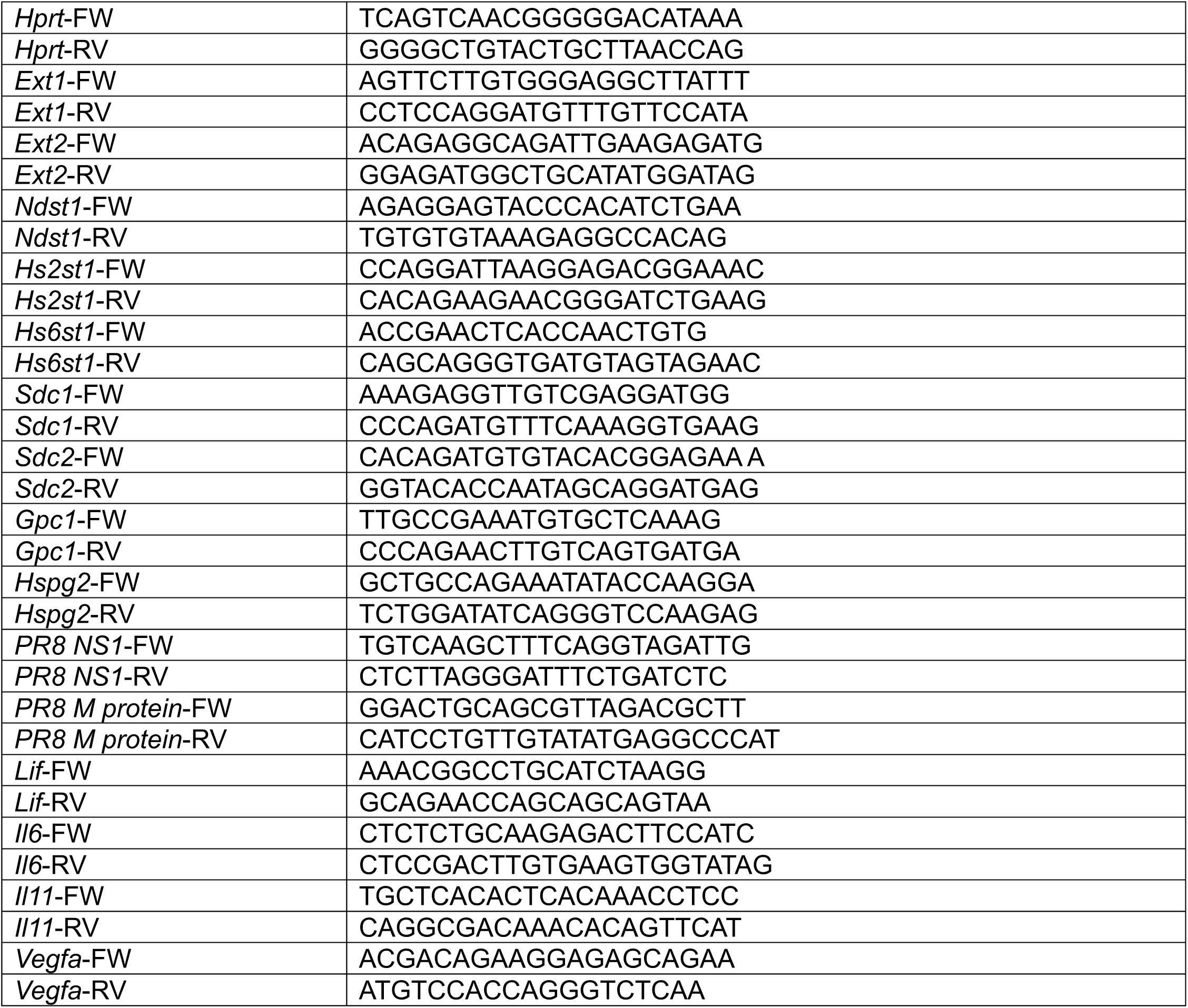
qPCR primers used in this study.

### Lung processing

Mice were euthanized and dissected to expose the lungs. Perfusion of the lungs was performed, after nicking the left femoral artery and the left atrium of the heart, through the left ventricle of the heart with 10 ml of cold 1x PBS. Bronchoalveolar lavage (BAL) was performed in the cases where tissue immune populations were analyzed or where bulk LMC was cultured directly after negative enrichment (this step was not done for sorting of Col14-LMC, sorting of Treg cells, or flow cytometric analysis of tissue mesenchymal cells). Lungs were extracted and placed in 0.5 ml tissue preparation media (RPMI with 100x penicillin/streptomycin, 100x GlutaMAX, 100x HEPES [all Gibco], and 5% FBS [Corning]) in a 5 ml Eppendorf tube, where they were minced. 3.5 ml was added to tubes of tissue preparation media with 5 U/ml DNAse, 1 mg/ml of collagenase A, and 1 mg/ml of dispase (pre-dissolved in 1x PBS consisting of 10% of the volume of the final digestion mixture). Lungs were digested in shaking incubator set to 110 r.p.m. at 37°C for 1h. Suspensions were then poured over a 100 µm cell strainer (Corning) into a 50 ml Falcon tube (Corning), pushed through the mesh with the top of a syringe, rinsed with 10 ml tissue preparation media, pushed through again, then rinsed with 5 ml tissue preparation media. Cells were centrifuged at 450 x g/4°C/5 min., then supernatants were poured off (due to delicate nature of pellet resulting from use of dispase). Pellets were resuspended in 2 ml of 1x ACK lysis buffer (deionized water with 154 mM ammonium chloride [Fisher], 10 mM potassium bicarbonate [Fisher], 0.1 mM ethylenediaminetetraacetic acid [EDTA] disodium salt dihydrate [Fisher], pH 7.2) and incubated at room temperature for 2 min., then quenched with 10 ml tissue preparation media and ran through a 100 µm nylon mesh sheet into a 15 ml Falcon tube (Corning). Cells were centrifuged at 450 x g/4°C/5 min., the supernatant was aspirated, and cells were resuspended in 1 ml of tissue preparation media and placed on ice. Cells were then used for antibody staining for flow cytometry or negative bead enrichment and/or sorting.

### Bulk LMC or CD4 T cell negative bead enrichment, and Col14-LMC and Treg sorting

Lungs were prepared as described in “Lung processing”. Lung cells were then negatively enriched for mesenchymal cells or CD4 T cells using iMag Streptavidin Particles Plus – DM beads (BD), according to their protocol. For bulk LMC and Col14-LMC isolation, biotinylated antibodies towards mouse CD45 (clone 30-F11; Biolegend), CD31 (clone 390; Biolegend), Epcam (clone G8.8; Biolegend), and TER-119 (clone TER-119; Biolegend) were used. For CD4 T cell isolation, biotinylated antibodies towards mouse CD31, Epcam, TER-119, Pdgfra (clone APA5; Biolegend), CD19 (clone 6D5; Biolegend), NK1.1 (clone PK136; Biolegend), CD11b (clone M1/70; Biolegend), CD11c (clone N418; Biolegend), and CD8a (clone 53-6.7; Biolegend) were used. For bulk LMC, following negative enrichment, cells were washed and resuspended in mesenchymal cell media (DMEM + 1x penicillin/streptomycin, 1x GlutaMAX, 1x sodium pyruvate, 1x nonessential amino acids [all Gibco], and 10% fetal bovine serum [FBS] [Corning]) and passed through 100 µm nylon mesh sheet to eliminate large conglomerates; cells were then washed again and resuspended in mesenchymal cell media for plating. Cells were confirmed to be <2% positive for CD45^+^, CD31^+^, or Epcam^+^ cells following enrichment. For Col14-LMC or Treg cell sorting, post-bead enrichment, cells were stained with fluorescent mouse antibodies for flow cytometric sorting (100 µl per mouse). For Col14-LMC sorting, antibodies used were: CD31-BV605 (clone 390; Biolegend), Epcam-PerCP-Cy5.5 (clone G8.8; Biolegend); Pdgfra-PE (clone APA5; Biolegend), CD146-PE-Cy7 (clone ME-9F1; Biolegend), CD45-APC (clone 30-F11; Cytek), and Sca-1-APC-Cy7 (clone D7; Biolegend). For Treg cell sorting, antibodies used were: CD45-BUV395 (clone 30-F11; BD), CD11b-BV510 (clone M1/70; BD), CD11c-BV510 (clone HL3; BD), TCR β-BV711 (clone H57-597; BD), CD45R (B220)-PerCP-Cy5.5 (clone RA3-6B2; Cytek), CD8a-PE (clone 53-6.7; Cytek), NK1.1-PE-Cy7 (clone PK136; Biolegend), and CD4-APC (clone RM4-5; Cytek). Treg cell preparations were done with lungs of *Foxp3*^EGFP^ mice, (i.e., Treg cells express GFP), allowing sorting for GFP^+^ cells. Cells were incubated with antibodies for 20 min. at 4°C. Post-staining, cells were washed, then resuspended in 200 µl per mouse, ran through a 70 µm mesh (for Col14-LMC sorting) or a 40 µm mesh (for Treg cell sorting), then sorted on a BD Aria sorter, with Sytox Blue added 5 min. prior to running samples for dead cell exclusion.

### Flow cytometry

Adherent cultured cells (LLC cells, A549 cells, bulk LMC, or Col14-LMC) were prepared for flow cytometry by washing with 1x PBS, then rinsing with 0.05% (LLC cells) or 0.25% (A549 cells, LMC) Trypsin-EDTA (Corning) for 5 min., following quenching with media. Ba/F3 cells (non-adherent) were directly removed from wells for flow cytometry. HS-modifying agents used in certain experiments are as described in “Western blotting” (without subsequent ligand treatment). For cell lines/cultured primary cells, antibody staining was performed with 25,000-250,000 cells, with ∼10,000 cells ran on a BD LSRII or BD Fortessa. Lungs were prepared for flow cytometry as described above, with 2-3 million cells from final single cell suspensions used for antibody staining and 100,000-1 million cells ran on a BD LSRII or BD Fortessa. Cells were stained in flow cytometry buffer (1x PBS with 1% BSA [Gold Biotechnology], 2.5 mM ethylenediaminetetraacetic acid [EDTA] disodium salt dihydrate [Fisher], and 0.1% sodium azide [Fisher]). Zombie Violet Fixable Viability Kit (Biolegend) or GhostDye Red 780 (Cytek), stained in a separate step from surface antibodies in 1x PBS), or Sytox Blue (Thermo), added directly to sample 5 min. prior to running on cytometer, were used for dead cell exclusion. For analysis of HS on cell lines or lung single cell suspensions, flow cytometry was done using HS-directed monoclonal mouse IgM antibodies 10E4 and/or JM403 (AMSBIO), followed by staining with Biotin-SP (long spacer) AffiniPure F(ab’)₂ Fragment Donkey Anti-Mouse IgM µ chain specific (Jackson ImmunoResearch), then staining with Streptavidin-APC (Invitrogen) or Streptavidin-Alexa Fluor 488 (Invitrogen); for mouse lung single cell suspensions, the Mouse-on-Mouse Immunodetection Kit (Vector Laboratories) was used for these staining steps. For wild-type LLC cells, HS core protein KO LLC cell sublines, and LMC, HS core proteins were assessed with flow cytometry using these anti-mouse antibodies: Sdc1-PE (clone 281-2; Biolegend), Sdc2 (polyclonal sheep IgG; R&D Systems), Sdc3 (polyclonal goat IgG; R&D Systems), Sdc4 (polyclonal rabbit IgG; Sigma-Aldrich), Gpc1 (polyclonal rabbit IgG, Invitrogen), Gpc3 (polyclonal rabbit IgG; Invitrogen), Gpc4 (polyclonal rabbit IgG; Proteintech), Gpc6 (polyclonal goat IgG; R&D Systems), and Hspg2 (rat monoclonal IgG, clone A7L6; Sigma-Aldrich). Live cells were used for all HS core protein stains, besides for Hspg2 staining where a Cytofix/Cytoperm Kit (BD) was used for fixation/permeabilization of cells. Besides for anti-Sdc1 which was conjugated to PE, cells were then stained with secondary antibodies Donkey Anti-Sheep IgG Alexa Fluor 647, Donkey Anti-Goat IgG Alexa Fluor 647, Donkey Anti-Rabbit IgG Alexa Fluor 647, Donkey Anti-Rat IgG Alexa Fluor 647 (all Invitrogen), or Donkey Anti-Rabbit Cy5 (Jackson Immunoresearch). For HS core protein analysis on LMC populations from lung single cell suspensions, cells were blocked with 10% normal horse serum (Gibco) in flow cytometry buffer, and stained with primary/secondary antibodies in 5% normal horse serum in flow cytometry buffer, prior to surface staining. For LMC analysis from lung single cell suspensions, surface staining was done using these anti-mouse antibodies: CD45-BUV395 (clone 30-F11; BD), Pdgfra-BV605 (clone APA5; Biolegend), CD31-BV711 (clone 390; Biolegend), Epcam-BV785 (clone G8.8; Biolegend); CD146-PerCPCy6.6 (clone ME-9F1; Biolegend), CD49e-PE (clone 5H10-27[MFR5]; Biolegend), Sca1-PE/Dazzle594 (clone D7; Biolegend), Pdpn-PE-Cy7 (clone 8.1.1; Biolegend), and CD9-APC-Fire750 (clone MZ3; Biolegend). Following surface marker staining, for staining with anti-mouse Ki67-AlexaFluor700 (clone SolA15; Invitrogen) or anti-mouse α-smooth muscle actin-eFluor660 (clone 1A4; Invitrogen), cells were fixed/permeabilized with Foxp3/Transcription Factor Staining Buffer Kit (Cytek). For lymphoid cell analysis from lung single cell suspensions, surface staining was done using these anti-mouse antibodies: NK1.1-BUV395 (clone PK136; BD), CD3-BUV496 (clone 145-2C11; BD), CD4-BUV737 (clone RM4-5; BD), TCR γ/δ-BV421 (clone GL3; Biolegend), CD11b-BV510 (clone M1/70; BD), CD11c-BV510 (clone HL3; BD), CD45-BV786 (clone 30-F11; BD), CD45R (B220)-PerCP-Cy5.5 (clone RA3-6B2; Cytek), CD8a-PE (clone 53-6.7; Cytek), TCR β-PE/Dazzle594 (clone H57-597; Biolegend), CD90.2 (Thy1.2)-PE-Cy7 (53-2.1; Biolegend); CD127 (IL-7Ra)-PE-Cy7 (A7R34; Cytek). Following surface staining, for staining lymphoid panel with anti-mouse Foxp3-FITC (clone FJK-16s; Invitrogen), cells were fixed/permeabilized with Foxp3/Transcription Factor Staining Buffer Kit (Cytek). For myeloid cell analysis from lung single cell suspensions, surface staining was done using these anti-mouse antibodies: CD45-BUV395 (clone 30-F11; BD), CD24-BV510 (clone M1/69; Biolegend), CD11b-BV650 (clone M1/70; Biolegend), CD103-BV711 (clone M290; BD), CD11c-FITC (clone N418; Cytek), Ly6C-PerCP-Cy5.5 (clone HK1.4; Biolegend), Siglec-F-PE (clone E50-2440; BD), Ly6G-PE/Dazzle594 (clone 1A8; Biolegend), CD64-PE-Cy7 (clone X54-5/7.1; Biolegend). Fluorescence-minus-one (FMO) controls were included to define staining boundaries where necessary.

### Ba/F3 electroporation

m*Egfr* sequence-containing pCMV6-Entry plasmid (Egfr [NM_207655] Mouse Tagged ORF Clone) was acquired from Origene (MR226160). The m*Egfr* sequence from the plasmid was amplified via PCR with addition of a stop codon at the end and flanking restriction enzyme cut sites (KpnI and NotI, beginning/end respectively), with resultant PCR product purified with Wizard SV Gel and PCR Clean-Up System (Promega). pCDNA3.0 plasmid was acquired from Invitrogen. pCDNA3.0 and amplified m*Egfr* PCR product (with additions) were cut with KpnI and NotI restriction enzymes (New England Biolabs), ran on an agarose gel, cut out of gel, and purified with Wizard SV Gel and PCR Clean-Up System (Promega). T4 DNA Ligase (New England Biolabs) was used to ligate cut plasmid and cut insert, with the product transformed into DH5α competent cells (Thermo). Individual clones were isolated on carbenicillin (Gold Biotechnology) agar plates, and Sanger sequencing (Azenta) was used to verify correct insertion (primers created by MacVector, ordered from Integrated DNA Technology). pCDNA3.0-m*Egfr* and pCDNA3.0 (empty vector) were transiently transfected into Ba/F3 cells using electroporation with a Gene Pulser Xcell Electroporation System (Bio-Rad) (25 μg uncut plasmid, 2.5 million Ba/F3 cells, 4 mm cuvette, 0.975 µF, 0.3 kV), then cultured with G418 (Sigma-Aldrich; 500 µg/ml) and rmIL-3 (Biolegend; 1 ng/ml) in Ba/F3 media (see “Cell Lines”). After outgrowth of electroporated cells in selection media, single clones were isolated by limiting dilution and, after outgrowth, assessed for EGFR protein expression by western blotting. A single clone derivative was used for rmEgfr and rmAreg treatment experiments.

### CRISPR-Cas9 knockout line generation in LLC cells

pSpCas9(BB)-2A-GFP (PX458) was acquired from Addgene (Plasmid # 48138). Single guide RNAs (sgRNAs) for selected mouse genes were generated using the Broad Institute sgRNA Designer. For each gene, two sgRNAs were selected targeting different exons. Selected sgRNAs were produced by Integrated DNA Technologies, with added overhangs for annealing into BbsI restriction enzyme (New England Biolabs)-cut PX458 (CACCG[sgRNA sequence] on forward annealing half and AAAC[sgRNA reverse complement sequence]C on reverse annealing half). sgRNAs were cloned into PX458 in a one-step reaction with BbsI and Quick Ligase (New England Biolabs), with products transformed into DH5α competent cells (Thermo). Individual clones were isolated on carbenicillin (Gold Biotechnology) agar plates, and Sanger sequencing (Azenta) was used to verify correct insertion (primers ordered from Integrated DNA Technology). Wild-type LLC cells were transfected with a mixture of Lipofectamine-2000 (Thermo) and both sgRNA-PX458 constructs (i.e., for both exons simultaneously) for each gene, diluted in Opti-MEM (Thermo). Media was changed after 6h, and 36h after transfection, cells were sorted on a BD Aria for GFP^+^ cells, into individual wells. Single cell clonal lines of cells were grown, and screened at the DNA level using specific primers for each exon targeted in each gene. Clones with DNA alteration were further analyzed at the RNA level (see “RNA extraction and qPCR”). Protein-level confirmation of KO or HS-level alterations conferred by KO was done with flow cytometry, as described above. For certain experiments, empty PX458 was transfected into LLC cells to create control cell lines.

### Transduction of LLC cells for KO rescue

MGC Mouse *Gpc1* cDNA (Horizon Discovery MMM1013-202859423) was acquired in a pYX-Asc vector. MSCV2.2 retroviral vector was acquired from Addgene (Plasmid #60206). Gibson assembly was used to clone *Gpc1* cDNA coding region into MSCV2.2 (strategy designed in MacVector). *Gpc1* was amplified from the original delivery vector, with primers including Gibson assembly overhangs, with Q5 High Fidelity 2X Master Mix (New England Biolabs), with resultant PCR product purified with Wizard SV Gel and PCR Clean-Up System (Promega). MSCV2.2 was cut with NotI (New England Biolabs), then ran on an agarose gel, cut out of gel, and purified with Wizard SV Gel and PCR Clean-Up System (Promega). Gibson Assembly Master Mix (New England Biolabs) was used with amplified *Gpc1* cDNA and cut MSCV2.2 to create MSCV2.2 with *Gpc1* insert (MSCV-*Gpc1*). Individual clones were isolated on carbenicillin (Gold Biotechnology) agar plates, and Sanger sequencing (Azenta) was used to verify correct insertion (primers created by MacVector, ordered from Integrated DNA technology). MSCV2.2 (MSCV-Empty vector [EV]) or MSCV-*Gpc1* were transfected into Phoenix-ECO cells with a mixture of plasmid and Lipofectamine-2000 (Thermo) diluted in Opti-MEM (Thermo). Transfected cells were passaged at 24h. At 48h post-transfection, viral supernatants were removed from wells, centrifuged at 450 x g/room temperature/5 min., and supernatants were ran through a 0.45 µm filter (Corning). Hexadimethrine bromide (polybrene) (5 ug/ml) Sigma-Aldrich) was added to filtered supernatants, then Gpc1-KO LLC cells were treated with viral supernatants. After ∼12h incubation, media was replaced, and transduced cells were passaged at 36h post-transfection. 48h later, transduced cells were sorted on a BD Aria for GFP^+^ cells (bulk sorted). Transduced cell lines were confirmed to maintain GFP and Gpc1 protein expression over several passages.

### siRNA treatment

ON-TARGETplus siRNA was acquired from Horizon Discovery/Dharmacon, with SMARTPools (4 siRNA constructs included) for mouse *Sdc2*, mouse *Gpc4*, or non-targeting siRNA control. Bulk LMC were isolated as described above and plated in mesenchymal cell media at 2 million cells/well in 6 well tissue culture-treated plates (Corning). Cells were allowed to adhere overnight (16-18h); the following day, media was aspirated from cells, cells were washed with DMEM (Gibco) to remove dead cells, and fresh mesenchymal cell media was added. Bulk LMC were then transfected with a mixture of siRNA and Lipofectamine RNAiMax Transfection Reagent (Thermo) diluted in Opti-MEM (Thermo). 48h post-treatment (no media change, siRNA left in wells throughout), cells were used for flow cytometry, or treated with rmAreg/rmEGF for western blotting (see above).

### Col14-LMC/Treg cell co-culture

Col14-LMC were isolated/sorted as described above and plated in mesenchymal cell media at 40,000-50,000 cells/well in 48 well tissue culture-treated plates (Corning). Cells were allowed to adhere overnight (16-18h); the following day, media was aspirated from cells, cells were washed with RPMI (Gibco) to remove dead cells, and fresh T cell media (RPMI + 100x penicillin/streptomycin, 100x GlutaMAX, 100x HEPES, 100x sodium pyruvate, 100x nonessential amino acids, 1000x β-mercaptoethanol [all Gibco], and 10% fetal bovine serum [FBS]) was added; cells were rested for ∼12h while Treg cell isolation/sorting occurred. Treg cells from lungs of IAV-infected mice (8 d.p.i.) were isolated/sorted as described above. 20,000-25,000 Treg cells (1:2 Treg cell:Col14-LMC ratio) were then added directly to Col14-LMC wells in small volumes (1:25 volume of media in wells). Dynabeads Mouse T-Activator CD3/CD28 for T-Cell Expansion and Activation (Thermo) beads were added to Treg cells at a 1:1 Treg cell/bead ratio prior to addition to wells. rhIL-2 (200 U/ml; NCI Preclinical Repository) and rhIL-7 (10 ng/ml; NCI Preclinical Repository) were added directly to wells at time of Treg cell addition. Cell co-cultures were incubated for 12h. At this time, wells were subjected to one wash of 5 mM EDTA in 1x PBS, and two additional washes of 1x PBS to remove Treg cells; following this removal, Col14-LMC in wells were lysed and analyzed for RNA (see “RNA extraction and qPCR”).

## References

1. Zaiss, D.M.W., et al., Emerging functions of amphiregulin in orchestrating immunity, inflammation, and tissue repair. Immunity, 2015. 42(2): p. 216–226.

2. Arpaia, N., et al., A Distinct Function of Regulatory T Cells in Tissue Protection. Cell, 2015. 162(5): p. 1078–89.

3. Kaiser, K.A., et al., Regulation of the alveolar regenerative niche by amphiregulin-producing regulatory T cells. J Exp Med, 2023. 220(3).

4. Xie, T., et al., Single-Cell Deconvolution of Fibroblast Heterogeneity in Mouse Pulmonary Fibrosis. Cell Rep, 2018. 22(13): p. 3625–3640.

5. Zepp, J.A., et al., Distinct Mesenchymal Lineages and Niches Promote Epithelial Self-Renewal and Myofibrogenesis in the Lung. Cell, 2017. 170(6): p. 1134–1148.e10.

6. Dahlgren, M.W., et al., Adventitial Stromal Cells Define Group 2 Innate Lymphoid Cell Tissue Niches. Immunity, 2019. 50(3): p. 707–722.e6.

7. Tsukui, T., et al., Collagen-producing lung cell atlas identifies multiple subsets with distinct localization and relevance to fibrosis. Nat Commun, 2020. 11(1): p. 1920.

8. Travaglini, K.J., et al., A molecular cell atlas of the human lung from single-cell RNA sequencing. Nature, 2020. 587(7835): p. 619-625.

9. Sarrazin, S., W.C. Lamanna, and J.D. Esko, Heparan sulfate proteoglycans. Cold Spring Harb Perspect Biol, 2011. 3(7).

10. Xu, D. and J.D. Esko, Demystifying heparan sulfate-protein interactions. Annu Rev Biochem, 2014. 83: p. 129–57.

11. Simon Davis, D.A. and C.R. Parish, Heparan sulfate: a ubiquitous glycosaminoglycan with multiple roles in immunity. Front Immunol, 2013. 4: p. 470.

12. Johnson, G.R. and L. Wong, Heparan sulfate is essential to amphiregulin-induced mitogenic signaling by the epidermal growth factor receptor. J Biol Chem, 1994. 269(43): p. 27149–54.

13. Cook, P.W., et al., Differential effects of a heparin antagonist (hexadimethrine) or chlorate on amphiregulin, basic fibroblast growth factor, and heparin-binding EGF-like growth factor activity. J Cell Physiol, 1995. 163(2): p. 418–29.

14. Nakanishi, H., et al., Structural differences between heparan sulphates of proteoglycan involved in the formation of basement membranes in vivo by Lewis-lung-carcinoma-derived cloned cells with different metastatic potentials. Biochem J, 1992. 288 (Pt 1)(Pt 1): p. 215-24.

15. Green, J.A., et al., A nonimmune function of T cells in promoting lung tumor progression. J Exp Med, 2017. 214(12): p. 3565–3575.

16. Schuksz, M., et al., Surfen, a small molecule antagonist of heparan sulfate. Proc Natl Acad Sci U S A, 2008. 105(35): p. 13075–80.

17. David, G., et al., Developmental changes in heparan sulfate expression: in situ detection with mAbs. J Cell Biol, 1992. 119(4): p. 961–75.

18. van den Born, J., et al., Presence of N-unsubstituted glucosamine units in native heparan sulfate revealed by a monoclonal antibody. J Biol Chem, 1995. 270(52): p. 31303–9.

19. Yarden, Y. and M.X. Sliwkowski, Untangling the ErbB signalling network. Nat Rev Mol Cell Biol, 2001. 2(2): p. 127–37.

20. Ornitz, D.M., et al., Receptor specificity of the fibroblast growth factor family. J Biol Chem, 1996. 271(25): p. 15292-7.

21. Collins, M.K., et al., Transfer of functional EGF receptors to an IL3-dependent cell line. J Cell Physiol, 1988. 137(2): p. 293–8.

22. Österholm, C., et al., Fibroblast EXT1-levels influence tumor cell proliferation and migration in composite spheroids. PLoS One, 2012. 7(7): p. e41334.

23. Zhou, P., et al., Epidermal growth factor receptor expression affects proliferation and apoptosis in non-small cell lung cancer cells via the extracellular signal-regulated kinase/microRNA 200a signaling pathway. Oncol Lett, 2018. 15(4): p. 5201–5207.

24. Ma, S., et al., The Hippo Pathway: Biology and Pathophysiology. Annu Rev Biochem, 2019. 88: p. 577–604.

25. Lampe, P.D. and D.W. Laird, Recent advances in connexin gap junction biology. Fac Rev, 2022. 11: p. 14.

26. Ryan, Z.C., et al., 1α,25-dihydroxyvitamin D(3) mitigates cancer cell mediated mitochondrial dysfunction in human skeletal muscle cells. Biochem Biophys Res Commun, 2018. 496(2): p. 746-752.

27. Zheng, B., et al., Ligand-dependent genetic recombination in fibroblasts: a potentially powerful technique for investigating gene function in fibrosis. Am J Pathol, 2002. 160(5): p. 1609–17.

28. Inatani, M., et al., Mammalian brain morphogenesis and midline axon guidance require heparan sulfate. Science, 2003. 302(5647): p. 1044-6.

29. Lucas, C.D., et al., Pannexin 1 drives efficient epithelial repair after tissue injury. Sci Immunol, 2022. 7(71): p. eabm4032.

30. Ito, M., et al., Brain regulatory T cells suppress astrogliosis and potentiate neurological recovery. Nature, 2019. 565(7738): p. 246-250.

31. Yayon, A., et al., Cell surface, heparin-like molecules are required for binding of basic fibroblast growth factor to its high affinity receptor. Cell, 1991. 64(4): p. 841–8.

32. Pye, D.A., et al., Regulation of FGF-1 mitogenic activity by heparan sulfate oligosaccharides is dependent on specific structural features: differential requirements for the modulation of FGF-1 and FGF-2. Glycobiology, 2000. 10(11): p. 1183–92.

33. Pellegrini, L., Role of heparan sulfate in fibroblast growth factor signalling: a structural view. Curr Opin Struct Biol, 2001. 11(5): p. 629–34.

34. Grzelak, E.M., et al., Pharmacological YAP activation promotes regenerative repair of cutaneous wounds. Proc Natl Acad Sci U S A, 2023. 120(28): p. e2305085120.

35. Prince, R.N., et al., The heparin-binding domain of HB-EGF mediates localization to sites of cell-cell contact and prevents HB-EGF proteolytic release. J Cell Sci, 2010. 123(Pt 13): p. 2308–18.

36. Lindahl, U., et al., Evidence for a 3-O-sulfated D-glucosamine residue in the antithrombin-binding sequence of heparin. Proc Natl Acad Sci U S A, 1980. 77(11): p. 6551–5.

37. Gutierrez, J. and E. Brandan, A novel mechanism of sequestering fibroblast growth factor 2 by glypican in lipid rafts, allowing skeletal muscle differentiation. Mol Cell Biol, 2010. 30(7): p. 1634–49.

38. Guéguinou, M., et al., *Lipid rafts,* KCa/ClCa/Ca2+ channel complexes and EGFR signaling: Novel targets to reduce tumor development by lipids? Biochim Biophys Acta, 2015. 1848(10 Pt B): p. 2603-20.

39. Colpitts, C.C. and L.M. Schang, A small molecule inhibits virion attachment to heparan sulfate- or sialic acid-containing glycans. J Virol, 2014. 88(14): p. 7806–17.

40. Clausen, T.M., et al., SARS-CoV-2 Infection Depends on Cellular Heparan Sulfate and ACE2. Cell, 2020. 183(4): p. 1043–1057.e15.

41. Mycroft-West, C.J., et al., Heparin Inhibits Cellular Invasion by SARS-CoV-2: Structural Dependence of the Interaction of the Spike S1 Receptor-Binding Domain with Heparin. Thromb Haemost, 2020. 120(12): p. 1700–1715.

42. Fontenot, J.D., et al., Regulatory T cell lineage specification by the forkhead transcription factor foxp3. Immunity, 2005. 22(3): p. 329–41.

43. Ge, S.X., E.W. Son, and R. Yao, iDEP: an integrated web application for differential expression and pathway analysis of RNA-Seq data. BMC Bioinformatics, 2018. 19(1): p. 534.

44. Reimand, J., et al., *g:Profiler-a web server for functional interpretation of gene lists (2016 update)*. Nucleic Acids Res, 2016. 44(W1): p. W83–9.

45. Stuart, T., et al., Comprehensive Integration of Single-Cell Data. Cell, 2019. 177(7): p. 1888–1902.e21.

46. De Vooght, V., et al., Oropharyngeal aspiration: an alternative route for challenging in a mouse model of chemical-induced asthma. Toxicology, 2009. 259(1-2): p. 84–9.

47. Pfaffl, M.W., A new mathematical model for relative quantification in real-time RT-PCR. Nucleic Acids Res, 2001. 29(9): p. e45.

